# Neonatal diethylstilbestrol exposure disrupts uterine epithelial apical-basal polarity and partial EMT state

**DOI:** 10.64898/2026.01.29.702663

**Authors:** Rachel E. Bainbridge, Wendy N. Jefferson, Tianyuan Wang, Sara A. Grimm, Carmen J. Williams

## Abstract

The developing female reproductive tract is highly sensitive to external hormonal stimulation, which can result in infertility and gynecologic diseases. To determine the underlying mechanisms, we used a mouse model to test the immediate, cell type-specific effects of neonatal exposure to the estrogenic chemical, diethylstilbestrol (DES), on the developing uterus. We found that control uterine epithelium is in a partial epithelial-mesenchymal transition state that is lost following DES exposure. This is accompanied by evidence of premature differentiation including altered apical-basal cell polarity and absence of the *Lgr5*+ epithelial stem cell population required for uterine gland formation. Cell-cell communication between epithelial and mesenchymal cells is restructured, and Wnt signaling is persistently reduced. The DES-exposed uterine mesenchyme has early signs of fibrosis through increased deposition of extracellular matrix collagen.Mechanistically, DES exposure causes cell type-specific changes in chromatin accessibility and gene expression, most prominently in epithelial cells. These changes can be explained in part by cell-specific alterations in chromatin looping at enhancer regions in concert with alterations in ERα binding. These findings suggest that reprogramming cell type-specific differentiation trajectories and extracellular matrix characteristics underlie the long-term phenotypic effects of developmental exposure to estrogenic endocrine disrupting chemicals. These changes lead to functional impairment of adult tissues and increased cancer risk.

**Significance Statement:** Uterine development is strongly impacted by brief exposure to estrogenic endocrine disruptors, but it is unclear why development is such a sensitive time point. This study employed multiomic analysis to identify cell type-specific uterine developmental trajectories in neonatal mice exposed to the estrogenic chemical, diethylstilbestrol, and compared these to controls. Control epithelium was under the influence of carefully orchestrated Wnt/β-catenin signaling and was in a partial epithelial-to-mesenchymal transition state. DES exposure repressed Wnt/β-catenin signaling and drove the epithelium toward full differentiation, resulting in the loss of both epithelial stem cells and normal apical-basal polarity. These changes provide an explanation for how endocrine disruptors can divert intrinsically programmed developmental trajectories to alter adult organ function.

## Introduction

Exposure to estrogenic endocrine disrupting chemicals during development causes an increased risk of infertility and cancer in women and in animal models (1). A classic example was the prenatal exposure of millions of women to the xenoestrogen, diethylstilbestrol (DES), which caused structural and functional reproductive tract anomalies and associated pregnancy complications. In addition, the exposed women are at higher risk for vaginal clear cell adenocarcinoma, uterine fibroids, endometriosis, and breast cancer (2). Despite confirmation in multiple animal models that endocrine disruption during development has distinct effects compared to adult responses to the same exposures, the reason(s) that development is a particularly sensitive window for endocrine disruption is not known.

We have been using a mouse model of exposure to the phytoestrogen genistein or to DES during postnatal reproductive tract differentiation to explore this question. Exposing neonatal mice to either of these estrogenic chemicals on postnatal days (PND) 1-5 results in infertility due in part to female reproductive tract dysfunction and development of uterine cancer in older adults (3-7). These phenotypes are accompanied by alterations in rostral-caudal patterning, a variable degree of failure of uterine gland development, and epithelial differentiation abnormalities (basal cell and squamous metaplasia) in addition to adenocarcinoma (8-12).

A partial epithelial-mesenchymal transition (EMT) that results in hybrid cell type states frequently occurs during tissue morphogenesis (13). This partial EMT is characteristic of processes as diverse as gastrulation and mammary gland branching morphogenesis; it is also observed in invasive and metastatic cancer cells (14). Cellular features of partial EMT vary widely but can include alterations in apical-basal polarity and cell adhesion to extracellular matrix (ECM) components. These changes are often accompanied by maintenance of classical epithelial features: stable epithelial junctional complexes linked to cytoskeletal elements that support inter-epithelial cell adhesion (14). Endometrial gland development follows a trajectory similar to that of mammary glands, with budding of epithelial cells into the underlying mesenchyme followed by cell proliferation and continued extension to create the final gland structures (15). In the mouse, this process begins around PND8 and requires expression of the Wnt signaling regulator, LGR5, which also serves as an epithelial stem cell marker (16). There is evidence in adult uterine tissue that estrogen exposure can repress *Lgr5* gene expression (17), but whether or not this occurs during neonatal uterine differentiation is unknown. In addition, cell type-specific impacts of DES or other estrogenic chemicals on uterine epithelial cell transition states are unknown.

Estrogen actions in the mouse uterus are largely mediated by estrogen receptor alpha (ERα) interactions with chromatin to modulate gene expression (18). ERα binds chromatin in the absence of ligand but changes its binding sites and binding partners following ligand activation (19). In adult mice undergoing regular estrous cycling, changes in steroid hormone levels across the cycle modulate ERα binding sites and 3-dimensional chromatin structure, along with changing chromatin accessibility and enhancer switching (20). In this setting, the progesterone receptor (PGR) is a key binding partner for ERα. How ERα influences gene expression in the neonatal mouse uterus and whether it interacts at that time of development with PGR is not known. On PND1, *Pgr* is not expressed except sporadically, but its expression rapidly increases through PND5, suggesting that PGR protein is likely present for at least some of this time and could influence ERα-mediated chromatin dynamics (21).

To explore the immediate effects of DES exposure on uterine cell identity, we performed single nucleus multiomic sequencing, which produces paired RNA-seq and ATAC-seq data from each cell, on PND5 uteri. We used these data to evaluate cell type-specific signaling interactions and validated some of these at the protein level. Our key findings were that the neonatal uterine epithelium is normally in a partial EMT state and that premature differentiation accompanied by alterations in apical-basal polarity and loss of epithelial stem cells are key cellular features explaining DES-induced adult uterine phenotypes. Identification of ERα binding sites and 3-dimensional chromatin structure in bulk PND5 uterine tissue provides a chromatin-centric view of how DES alters cell differentiation trajectories to interfere with proper uterine development and growth.

## Results

### DES exposure induces gene expression changes largely in epithelial and mesenchymal cells

To explore cell-type specific impacts of neonatal DES exposure on gene expression and chromatin accessibility in early uterine development, we performed single-nucleus multiomic sequencing of the uterus of DES-exposed or control mice on PND5. From control uteri, 20,000 cells (max cutoff) were obtained with 2,571 median genes per cell and 1,229 high-quality ATAC fragments per cell. From DES-exposed uteri, 13,164 cells were obtained with 1,473 median genes per cell and 3,335 high-quality ATAC fragments per cell. Following initial quality control, 13,154 cells from control uteri and 7,339 cells from DES-exposed uteri were used for analysis.

To determine differences in uterine cell type composition, we assessed cell identity in an integrated dataset of both DES-exposed and control cells using the single nucleus RNA sequencing (snRNAseq) data. An integrated UMAP (uniform manifold approximation and projection) of all captured uterine cells showed 11 distinct clusters (Figure 1A; Dataset S1). We performed manual cell identity annotation using markers published in a single cell RNAseq (scRNAseq) dataset from C57BL/6J uteri at PND3 (22) (Figure S1A). Cells clustered mainly by cell identity (Figure 1B). There were no obvious differences in cell clusters between exposure groups except for the epithelial cells, which were generally non-overlapping between control and DES (Figure 1C). Though there were fewer cells within each DES-exposed cell identity group, the relative proportions of each were similar between DES-exposed and control uteri, except in myocytes and endothelium (Figure S1B). Gene expression across cell types and in the two exposure groups was compared by generating a heatmap of the top 25 differentially expressed genes per cell type and exposure (Figure 1D, Dataset S2). DES-exposed epithelial and mesenchymal cells had overall lower expression of the control markers and also expressed unique marker genes. This phenomenon was not observed in the remaining cell types, which had similar levels of marker gene expression in both exposure groups.

**Figure 1.**
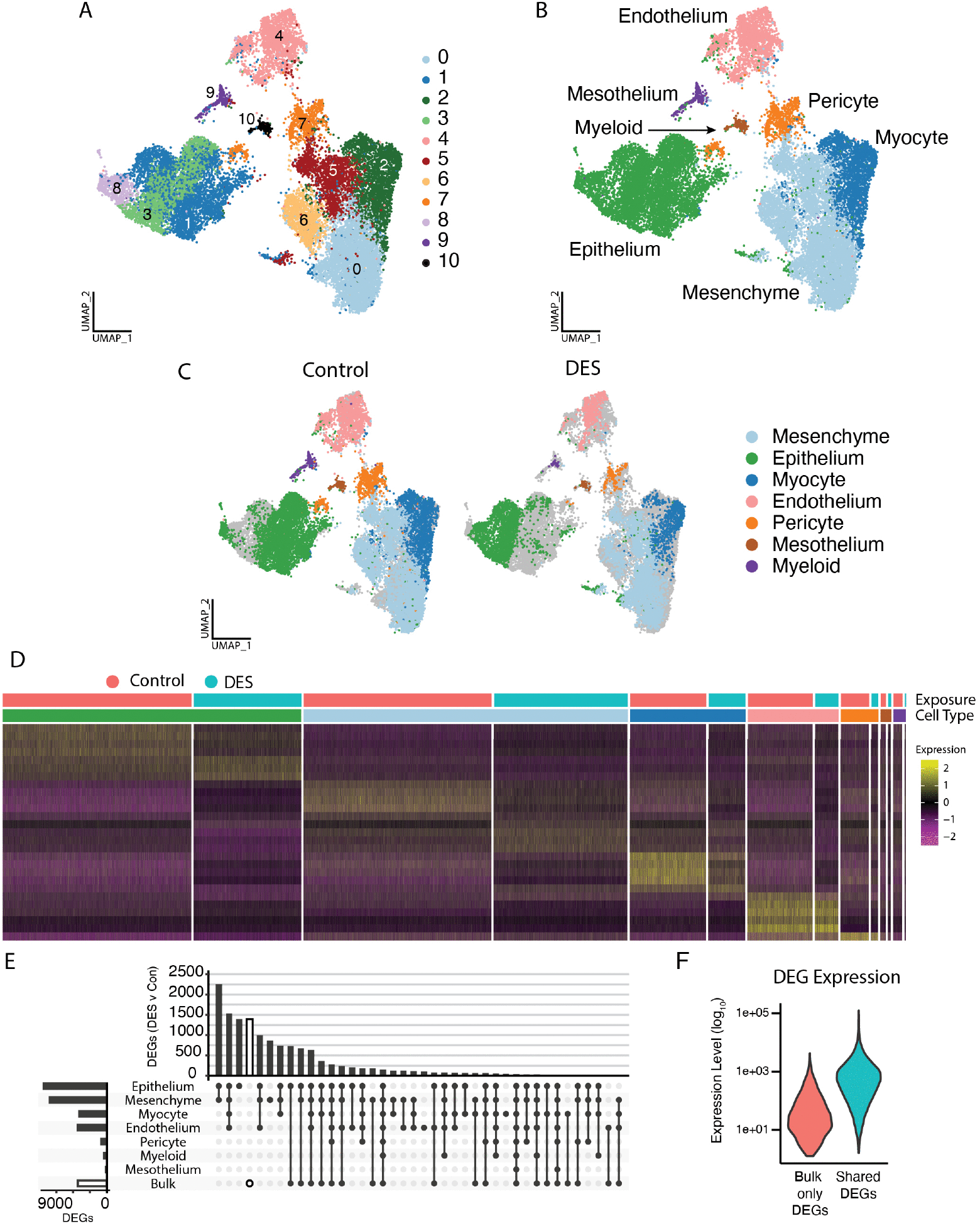
Single-nucleus RNAseq demonstrates differences in gene expression between control and DES-exposed uterine mesenchyme and epithelium. A) Integrated UMAP of all cells captured with single nucleus RNAseq from control or DES-exposed uteri labeled by cluster. B) Integrated UMAP with manually annotated cells including mesenchyme, epithelium, endothelium, stroma, pericyte, mesothelium, myocyte, and myeloid cells. (See Figure S1) C) Annotated UMAP split by treatment. D) Heatmap of RNA expression across all cells of top marker genes for each cell type per exposure. E) UpSet plot of differentially expressed genes between DES-exposed and control cells in each cell type or previously published bulk sequencing (23). F) Expression level of differentially expressed genes appearing only in bulk RNAseq vs those appearing both in bulk and snRNAseq (shared).

Differentially expressed genes (DEGs) between DES and control were determined for each cell type (Dataset S2; 18,038 genes across all cell types). These were compared to the list of DEGs from our published PND5 bulk uterine RNAseq dataset from mice exposed to DES under the same protocol (23) (5,199 genes). Many DEGs were identified by snRNAseq in one or more of the uterine cell types that were not identified in the bulk RNAseq (9208/13013, 70.8%) (Figure 1E). In addition, many genes also were uniquely differentially expressed in the bulk RNAseq that were not captured by snRNAseq (1394/5199, 26.8% of bulk DEGs; Figure 1E); most of these DEGs exhibited lower gene expression than genes that overlapped snRNAseq DEGs (Figure 1F). These observations demonstrate the differential utility of both approaches, with cell specificity provided by one approach and greater ability to detect less abundant RNAs by the other. The majority of the snRNAseq DEGs were differentially expressed in both epithelium and mesenchyme, with about half of these also DEGs in endothelium.

To explore DES-induced changes in gene expression in epithelial and mesenchymal cells, we restricted the integrated whole uterine dataset to only epithelial cells and mesenchymal cells. Because myocytes clustered closely with mesenchymal cells, with some myocytes overlapping the mesenchymal cluster, myocytes were also included with mesenchymal cells. In the resulting UMAP, clusters were based almost entirely on exposure status, masking any cell type-specific differences between populations (Figure S2A). For this reason, the epithelial/mesenchymal/myocyte dataset was split by exposure status and each exposure group analyzed separately (Figure S2B-C). In both control and DES UMAPs, cells were defined largely by epithelial or mesenchymal characteristics (Figure S2D-G; Datasets S3-S4). Within the mesenchymal population, though cells most strongly expressed mesenchymal markers, they also expressed either stromal (*Vcan, Dcn*) or myocyte (*Myh11, Chrm3*) markers at a lower level (11, 22, 24). Epithelial and mesenchymal (stroma-like, but not myocyte-like) populations were subset from these datasets and used in subsequent analyses to explore differences within each cell type due to DES exposure. Both control and DES-exposed epithelial cells (Figure 2A) and control and DES-exposed mesenchymal cells (Figure 2B) formed discrete clusters (epithelial, 6 clusters; mesenchymal, 5 clusters). DES exposure resulted in 906 upregulated and 3,090 downregulated genes in epithelial cells (Dataset S5) and 774 upregulated and 930 downregulated in mesenchymal cells (Dataset S6), which were further explored.

**Figure 2.**
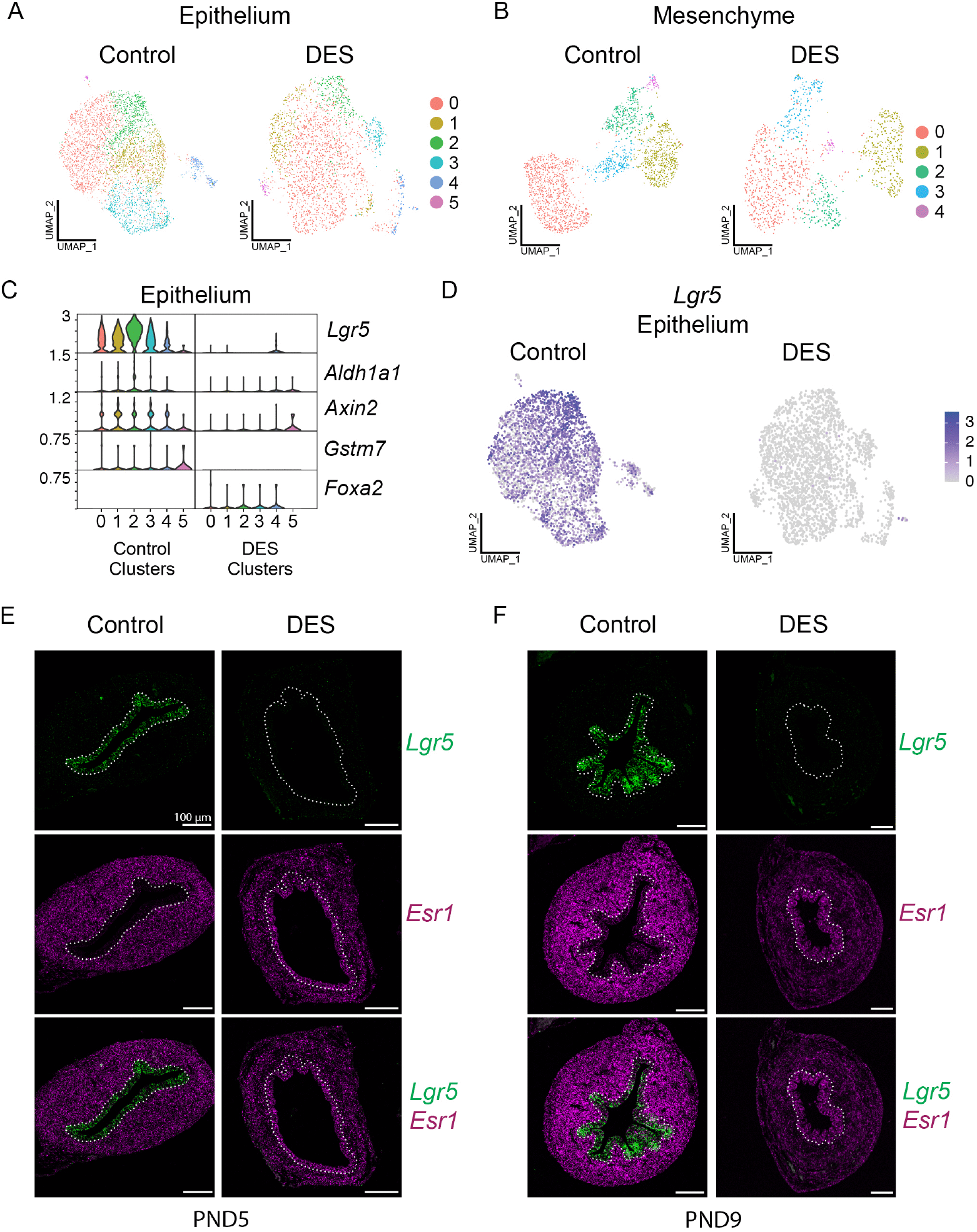
DES-exposed epithelium does not have typical stem cell populations. A) UMAPs of cells from control or DES-exposed epithelium. B) UMAPs of cells from control or DES-exposed mesenchyme. C) Violin plots of stem cell marker genes *Lgr5, Aldh1a1, Axin2, Gstm7* and mature gland marker gene *Foxa2* in control or DES-exposed epithelium. D) Feature maps showing expression of *Lgr5* in control or DES-exposed epithelium. E-F) Representative images of cross sections of PND5 (E) or PND9 (F) uterine horns from control or DES-exposed mice, with fluorescent RNA probes for *Lgr5* (top, green) and *Esr1* RNA (pink, middle; overlay, bottom). The epithelium is outlined with dotted lines. PND5 Control, N = 15 from 3-7 mice; PND5 DES, N = 10 from 3-6 mice; PND9 Control, N = 9 from 4 mice; PND9 DES, N = 4 from 4 mice.

### PND5 DES-exposed epithelial cells lack a typical stem cell population

We demonstrated previously that epithelial stem cells could not be identified in the uterus of 12-month-old adults exposed neonatally to DES (11). To determine if this population was lost with age or was never properly established, we examined markers of stem cell populations at PND5. Control PND5 epithelial cells expressed classic stem cell markers, including *Lgr5, Aldh1a1, Axin2*, and *Gstm7*; *Lgr5* exhibited the highest expression (Figure 2C) (11). The highest expression of *Lgr5* was observed in control cluster 2, but it was also expressed in most cells of all other clusters (Figure 2C-D). The stem cell markers were minimally or not expressed in DES-exposed epithelium. However, a marker of mature glandular epithelium, *Foxa2*, was expressed in DES-exposed epithelium but absent in control epithelium, suggesting that DES induces premature differentiation of the uterine epithelium.

To validate the *Lgr5* expression using a second methodology, we performed in situ hybridization on cross-sections of uterine tissue. *Esr1* was used as a positive control for the method as the protein is detected in the neonatal female reproductive tract (25, 26). In PND5 controls, *Lgr5* was detected in most epithelial but not mesenchymal cells, while *Esr1* was detected only in mesenchymal cells (Figure 2E). In DES-exposed epithelium, *Lgr5* was not detected but *Esr1* was expressed in both epithelium and mesenchyme. These findings are consistent with previous reports of loss of uterine *Lgr5* expression following estradiol treatment of ovariectomized adults (17). To determine if the DES-induced loss of *Lgr5* expression persisted to the time when gland formation begins, we performed in situ hybridization on control and DES-exposed uteri at PND9. *Lgr5* was readily detected in control epithelium but not in DES-exposed uteri (Figure 2F). *Esr1* was minimally expressed in the epithelium and highly expressed in the mesenchyme in control uteri. DES-exposed uteri continued to have *Esr1* expression in epithelial cells and reduced levels in mesenchyme compared to controls on PND9 (similar levels to PND5 control mesenchyme) suggesting failure to up-regulate *Esr1* in DES-exposed mesenchyme. These data support the idea that neonatal DES exposure induces premature epithelial cell differentiation and loss of the existing stem cell population after only 5 days of treatment.

### DES-exposure alters cell-cell communication in the developing uterus

The communication between mesenchyme and epithelium is vital for proper reproductive tract development (25, 27, 28). To explore the global capacity of these cell types to communicate following DES exposure, we assessed expression of known ligand-receptor pairs using the R package, LIANA (29). The relative number of all ligand-receptor pairs in both epithelium and mesenchyme in control and DES uteri is shown as a chord map (Figure 3A). Control and DES cells had similar percentages of mesenchyme-to-mesenchyme signal pairs but differed in all other signal pair patterns. Mesenchyme to epithelium signaling predominated in control cells, while epithelial to mesenchyme signaling predominated in DES cells. Epithelium-to-epithelium signaling pairs were present in controls, but the small number of such pairs in DES cells was insufficient to visualize in the chord map.

**Figure 3.**
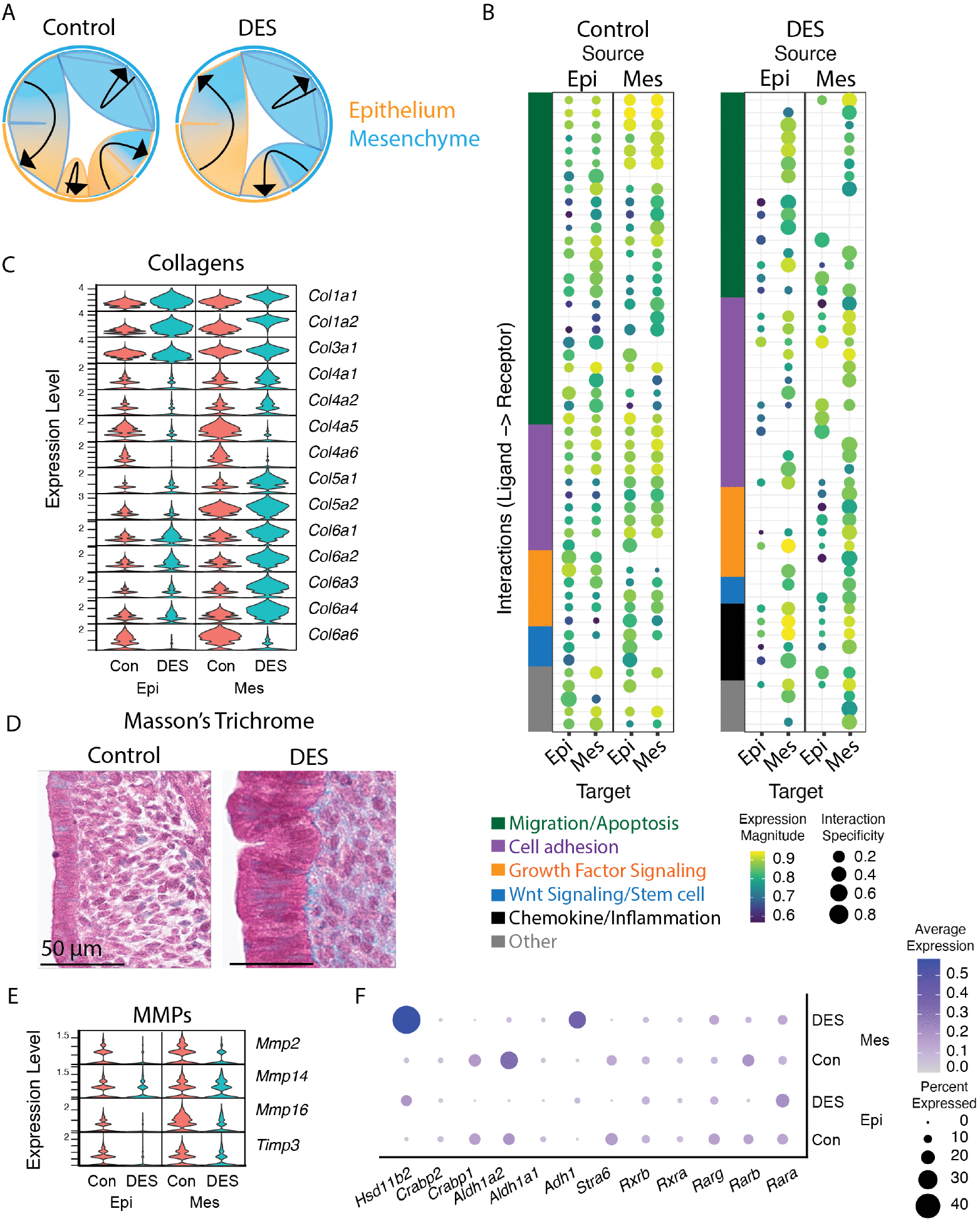
DES exposure alters uterine epithelial-mesenchymal communication. A) Chord map depicting relative abundance of possible ligand-receptor pairs between mesenchyme-mesenchyme (blue), mesenchyme-epithelium (blue to orange), epithelium-mesenchyme (orange to blue), and epithelium-epithelium (orange). Arrow directions are from source to target. B) Dot plots representing the top 50 ligand-receptor pairs available to interact (by presence of RNA) in control or DES-exposed epithelium or mesenchyme. Label at top notes ligand source; label at bottom notes target receptor source. Each row represents a ligand-receptor pair, clustered and color coded to denote function. Ligand-receptor pairs included are listed in Dataset S7. C) Violin plots of RNA expression of collagen genes expressed in control or DES-exposed epithelium or mesenchyme. D) Representative Masson’s trichrome staining of cross sections of PND5 uterine horn from control or DES-exposed mice (blue denotes collagen; Control, N = 11 from 7 mice; DES, N = 12 from 6 mice). E) Violin plots of RNA expression of matrix metalloprotease genes in control or DES-exposed epithelium or mesenchyme. F) Dot plot of retinoic acid signaling gene expression in control or DES-exposed epithelium and mesenchyme.

To examine the functions of the predicted intercellular signaling pairs, we plotted the top 50 ligand-receptor pairs from each condition by rank aggregate score (Figure 3B; Datasets S7, S8,S9). These ligand-receptor pairs were grouped by function to reveal patterns in functional gains or losses in communication. In controls, migration and/or apoptotic signaling pathways had the largest number of ligand-receptor pairs. [Note that migration and apoptosis are grouped in this analysis as many of these pairs act in shared pathways that balance the two processes (30)]. Additional pathways well represented in controls were growth factor signaling and cell adhesion. The pairs represented in these three pathways generally had ligands and receptors in both epithelium and mesenchyme, suggesting active communication within and between both cell types. Control cells had only 3 ligand receptor pairs in the Wnt signaling pathway, and these were different in that the ligands were expressed in both epithelium and mesenchyme, but the receptors were only in the epithelium. This finding indicates that Wnt signaling in controls is directed toward epithelial cell responses. Pairs annotated as “other” had assorted functions that did not fit into a larger group with shared function.

In DES cells, the most well represented pathways were similar to controls, however, there were far fewer pairs in migration/apoptosis and far more in cell adhesion pathways (Figure 3B). In addition, rather than ubiquitous cell type communication predicted in controls, there was minimal predicted epithelium-to-epithelium communication in DES cells (Figure 3A). The essential Wnt signaling pathway was represented by two ligand-receptor pairs. However, there was no epithelial to epithelial Wnt signaling and the critical mesenchyme to epithelium signaling was reduced or absent. In addition, a distinct signaling pathway, chemokine/inflammation, was represented by six ligand-receptor pairs in the DES cells but none of these were observed in controls. Overall, DES-exposed uteri contained epithelial cells that were less communicative with each other, less receptive to mesenchymal signals, and most likely receive more adhesive and inflammatory signals and fewer growth and migratory signals than controls. This finding is consistent with previous observations of a dramatic reduction in cell proliferation following DES exposure (8, 9, 31).

Many of the predicted upregulated cell adhesion ligands were collagens, which are key extracellular matrix components. The most highly expressed collagens were fibril-forming types I, III and V, with two type I isoforms (*Col1a1* and *Col1a2*) and two type V isoforms (*Col5a1* and *Col5a2*) highly upregulated following DES exposure in mesenchyme and the type I isoforms were also upregulated in epithelium (Figure 3C). Genes encoding type IV collagen, a major component of basement membranes, were generally downregulated following DES exposure in both epithelium and mesenchyme. Multiple type VI collagen isoforms, which form beaded filaments rather than fibrils, were also highly upregulated in the mesenchyme following DES exposure. Consistent with these observations, Masson’s trichrome staining revealed that collagen was excessively accumulated in DES uterine mesenchyme compared to controls (Figure 3D). There was also very intense red staining of the DES epithelial cells, suggesting increased levels of keratins. This finding was consistent with the dramatically increased expression of *Krt7, Krt8*, and *Krt19* in DES epithelium (Dataset S5). Matrix metalloproteases degrade collagen in mesenchyme, which supports epithelial invasion during gland development (32-34). In controls, genes encoding matrix metalloproteases (*Mmp2, Mmp14*, and *Mmp16*) were expressed in epithelium and at a higher level in mesenchyme; these changes were accompanied by expression of tissue inhibitor of metalloprotease 3 (*Timp3*) (Figure 3E). Expression of all of these genes was decreased with DES exposure, suggesting that both increased collagen deposition and reduced matrix metalloprotease expression contribute to the palpably increased stiffness of the DES-exposed uterus at PND5.

Retinoic acid signaling is implicated in the mesenchymal-epithelial crosstalk that orchestrates development of the female reproductive tract. In adult mice, retinoic acid signaling limits the impact of estrogen signaling on the uterine epithelial phenotype, helping to maintain it as a simple columnar rather than stratified epithelium (35). There is also evidence for the converse, that estrogen signaling in the uterus may limit retinoic acid signaling (35). In the current study, retinoic acid signaling mediators were not identified as ligand-receptor pairs in either control or DES uteri (Datasets S10, S11). Although both epithelial and mesenchymal cells expressed some retinoic acid signaling receptors, mediators, and targets, these were detected in very few cells and at a low expression level in either compartment (Figure 3F). An exception was *Hsd11b2*, a retinoic acid signaling target that was upregulated, rather than downregulated, in the mesenchyme following DES exposure. These findings suggest that the PND5 uterus is not yet relying on retinoic acid signaling to limit estrogen effects, consistent with the lack of endogenous estrogen at this time in development.

### DES exposure suppresses Wnt signaling

The Wnt/β-catenin signaling pathway is important for regulating growth and development, with nuclear translocation of β-catenin and interaction with TCF/LEF transcription factors key for its transcriptional activity (36, 37). Following neonatal DES exposure, expression of Wnt signaling ligands, receptors, and targets is dramatically increased in uterine epithelial cells at 12 months of age (11), but it was unknown when this increased activity began. Neonatal DES exposure suppresses some Wnt signaling mediators (*Wnt7a, Wnt11, Wnt16*, and *Fzd10*) in the PND5 uterus as indicated by in situ hybridization or microarray methods (8). To determine at a cell type-specific level how Wnt signaling was likely to be impacted immediately following neonatal DES exposure, we generated violin plots of the expressed genes that participate in regulating Wnt signaling at the plasma membrane (Figure 4A). In PND5 control epithelium, the most strongly expressed components of signaling directly at the frizzled receptor complex were the activating ligands, *Wnt5a* and *Wnt7a*, the inhibitory ligand, *Sfrp2*, and transmembrane components, *Fzd3, Fzd6*, and *Lrp6*. Components of the plasma membrane complex responsible for ubiquitinating frizzled receptors were also expressed strongly, including *Lgr5* and *Znrf3. Rspo* genes, which encode the ligands that promote downregulation of the LGR/ZNRF3 complex, were not expressed in the epithelium. Control mesenchyme expressed additional secreted regulators that could impact Wnt signaling in the epithelium, including both activators (*Wnt4, Wnt5a*, and *Wnt5b*), and inhibitors (*Dkk2* and *Sfrp2*). Overall, this gene expression profile provided a picture of balanced Wnt signaling activation that was tempered by both ligand-based inhibition and proteolytic degradation. The control mesenchyme had similarly balanced Wnt signaling gene expression, with *Fzd2, Fzd3*, and *Lrp6* expressed and potentially activated by several expressed Wnts. Proteolytic degradation to counteract this activation would be expected through strong *Znrf3* expression because the absence of *Lgr* gene expression to produce an LGR receptor would prevent RSPO3 from acting to turn over ZNRF3.

**Figure 4.**
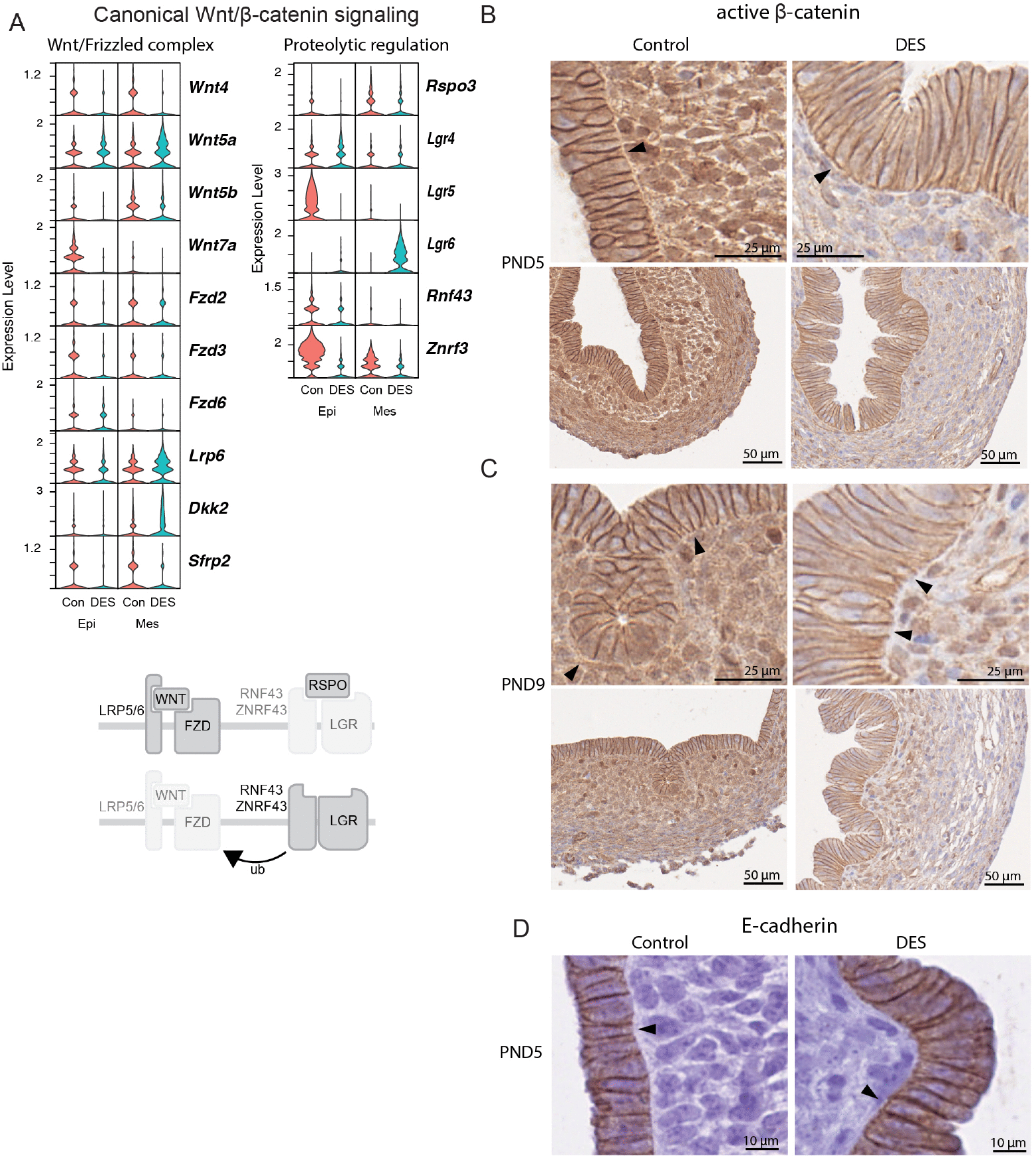
DES exposure suppresses Wnt/β-catenin signaling. A) Violin plots of Wnt/μ-catenin complex members in control or DES epithelium or mesenchyme. Diagram demonstrates control of Wnt signaling by LGR and RNF43/ZNRF43 through ubiquitination of the Frizzled/LRP complex. B-C) Representative images of active (non-phosphorylated) μ-catenin immunohistochemical staining in control or DES-exposed uterus at B) PND5 (Control, N = 4 from 4 mice; DES, N = 6 from 6 mice) or C) PND9 (Control, N = 8 from 8 mice; DES, N = 7 from 7 mice). D) Representative images of E-cadherin immunohistochemistry in control or DES-exposed uterus at PND5 (Control, N = 14 from 3-6 mice; DES, N = 12 from 3-6 mice).

In PND5 DES uteri, *Wnt4* and *Wnt7a* were downregulated in the epithelium and *Wnt5a* was upregulated in mesenchyme (Figure 4A). There was negligible expression of any *Fzd* genes in either epithelium or mesenchyme. In addition, genes encoding components of the proteolytic regulation complex were expressed, including *Lgr4* in the epithelium, *Lgr6* in the mesenchyme, and *Znrf3* in both. These findings suggested a complete lack of Wnt signaling immediately following DES exposure.

To determine if these suggested differences in Wnt signaling were reflected at the protein level, we examined PND5 uteri for the presence of nuclear active (non-phosphorylated) β-catenin as an indicator of β-catenin-driven transcription. Controls had active β-catenin immunostaining in many but not all mesenchymal and epithelial nuclei, though nuclear epithelial signal was difficult to ascertain with confidence due to cellular crowding and the presence of intense β-catenin staining at the cell-cell borders (Figure 4B). In contrast, DES uteri had very few mesenchymal nuclei with active β-catenin immunostaining and nuclear immunostaining in the epithelium was likely absent although crowding precluded a definitive answer. These findings were consistent with the gene expression profiles suggesting balanced Wnt signaling in controls but suppression of Wnt signaling in DES-exposed uteri. In PND9 DES-exposed uteri, the peri-epithelial mesenchyme had some nuclear immunostaining while the remainder of the mesenchyme and epithelium did not (Figure 4C), indicating partial recovery of mesenchymal but not epithelial Wnt signaling activity soon after discontinuing DES.

### DES exposure alters epithelial apical-basal polarity and partial EMT state

In addition to its transcriptional activity, β-catenin has an important role in supporting the cytoskeleton that is critical for integrity of epithelial monolayers. At bicellular adherens junctions, β-catenin bridges extracellular connections between cadherin molecules to the intracellular actin cytoskeleton through binding both the E-cadherin C-terminus and α-catenin (38). Consistent with this role in cytoskeletal regulation, at PND5 there was strong non-phosphorylated β-catenin staining at the plasma membrane of epithelial cells in both treatment groups (Figure 4B). A subtle difference between groups was that controls had β-catenin signal only at the lateral plasma membrane region whereas DES-exposed uteri also had β-catenin at the basal surface, adjacent to the basement membrane (Figure 4B). E-cadherin localized to the same sites as β-catenin in both control and DES-exposed uteri, though the E-cadherin basal plasma membrane staining in DES-exposed uteri was more punctate than β-catenin (Figure 4D). Furthermore, the active β-catenin and E-cadherin staining in control PND5 uteri revealed a clearly delineated basal lamina that was difficult to visualize in DES-exposed uteri (Figure 4B, 4D). Taken together with the expanded number of ligand-receptor pairs related to cell adhesion in the DES-exposed uterus (Figure 3B), these findings suggest that at PND5 the two groups have differences in epithelial adhesion characteristics.

Bidirectional signaling between ECM molecules and epithelial cells is vital for the initial establishment and maintenance of apical-basal cell orientation and cell adhesion to the underlying basement membrane. Key mediators of this communication include ECM collagens and laminins. Many collagen genes were highly expressed in the PND5 uterus, particularly following DES exposure (Figure 3C, 3D). All laminins are heterotrimers composed of α, β, and γ subunits, with the α subunit physically interacting with integrins. In control epithelium, four laminin α subunits (*Lama2, Lama3, Lama4*, and *Lama5*) were expressed at very low levels, whereas the DES epithelium only expressed *Lama5* and did so much more strongly than in controls (Figure 5A). In the mesenchyme, the most highly expressed laminins were *Lama2* and *Lama4*, with *Lama4* downregulated following DES. Of the genes encoding laminin beta and gamma chains, *Lamb1* and *Lamc1* were most highly expressed. Together, these findings suggest that excess laminin-511 (α5β1γ1) was generated by DES epithelium, while laminin-211 and laminin-411 were produced mainly in mesenchyme of both treatment groups.

**Figure 5.**
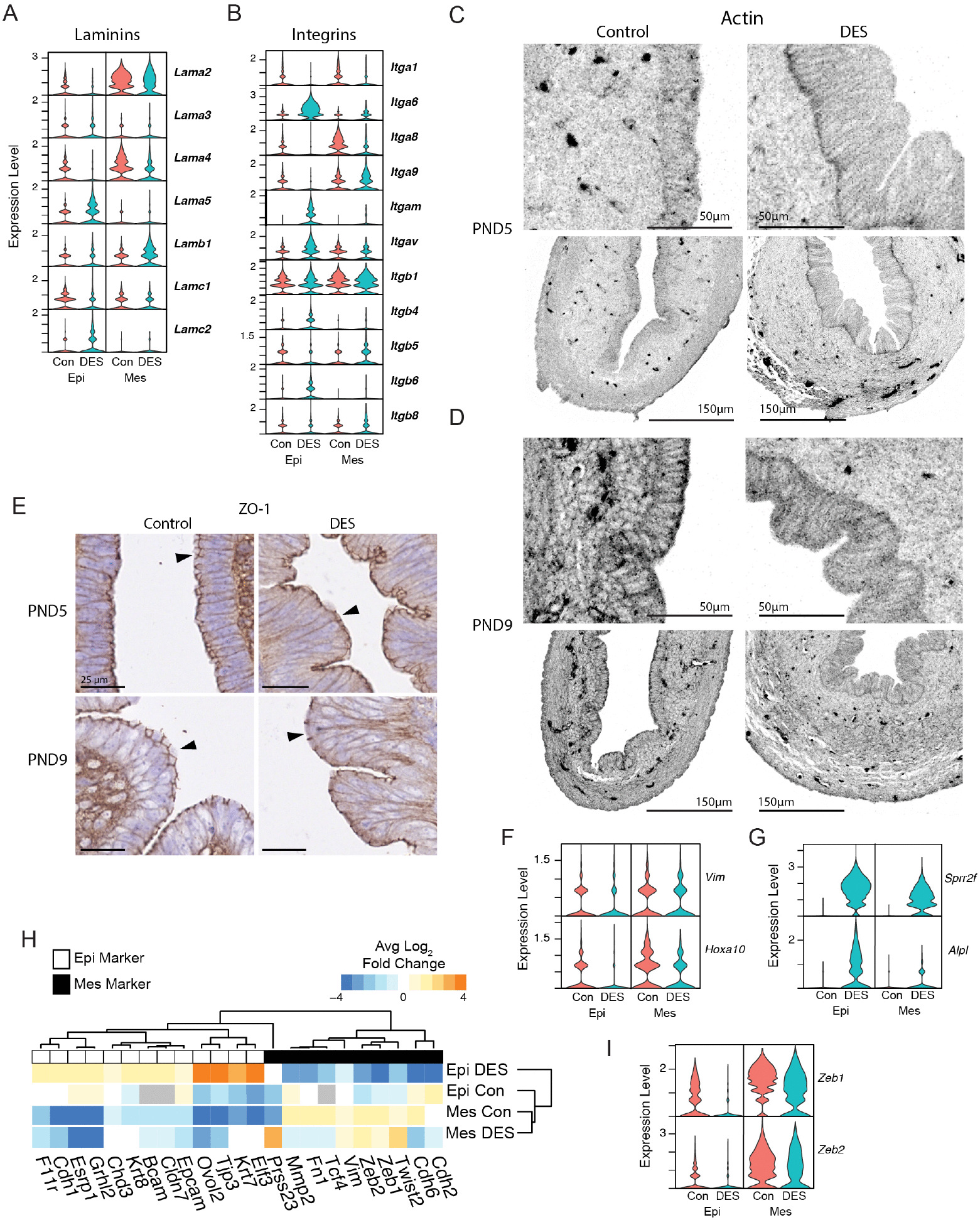
DES exposure alters epithelial cell polarity and EMT state. A) Violin plots of RNA expression of laminin genes in control and DES-exposed epithelium and mesenchyme. B) Violin plots of RNA expression of integrin genes in control and DES-exposed epithelium and mesenchyme. C-D) Representative images of filamentous actin staining in C) PND5 or D) PND9 control or DES-exposed uterus (Control, N = 7 from 5 mice; DES, N = 7 from 5 mice). E) Representative images of immunohistochemical staining of tight junction protein ZO-1 in control and DES-exposed uterus at PND5 (Control N = 8 from 5 mice, DES N = 9 from 5 mice). F) Violin plots of RNA expression of mesenchyme marker genes *Vim* and *Hoxa10* in control or DES-exposed epithelium or mesenchyme. G) Violin plots of RNA expression of terminal epithelial differentiation genes, *Sprr2f* and *Alpl*, in control or DES-exposed epithelium or mesenchyme. H) Hierarchical heatmap showing fold change of RNA expression of EMT marker genes in control or DES-exposed epithelium or mesenchyme. Black or white boxes along top axis represent whether gene is an epithelium (Epi) or mesenchymal (Mes) marker. I) Violin plots of RNA expression of EMT-controlling transcription factors, *Zeb1* and *Zeb2*, in control or DES-exposed epithelium or mesenchyme.

Adhesion of epithelial cells to the underlying basement membrane relies largely on the epithelial cell expression of the integrin heterodimers that serve as laminin receptors: integrins α3β1, α6β1, α7β1, and α6β4 (39). *Itgb1* was strongly expressed in both epithelial and mesenchymal cells regardless of treatment group, but *Itgb4* was only expressed at a low level following DES exposure, not in controls (Figure 5B). In control epithelial cells, *Itga6* was expressed at a very low level, but it was highly upregulated in epithelium following DES exposure; neither group expressed *Itga3* or *Itga7*. These findings suggest that DES epithelial cells likely expressed mainly integrins α6β1 and α6β4, appropriate laminin receptors to cause increased adhesiveness to the underlying basement membrane.

Epithelial sheets maintain their structure through stabilizing actomyosin cytoskeletal networks in a belt near the apical region (40). Developing intestinal epithelium has an additional actomyosin network at the basal region, except at regions where crypts will form by protrusion of the epithelium into the underlying mesenchyme (41). To determine if the changes we observed in collagen, laminin, and integrin expression were correlated with alterations in establishment of epithelial actomyosin skeletal networks in the uterus, we examined localization of filamentous actin. In control PND5 epithelial cells, actin staining was visible in a diffuse pattern at the apical region and in puncta at cell-cell junctions (Figure 5C). In comparison, DES PND5 epithelium had similar puncta at cell-cell junctions but had strong actin staining at the basal region adjacent to the basement membrane and minimal apical actin. By PND9, controls had increased overall actin staining, with higher intensity staining at the apical region and some regions that had clear basal actin staining (Figure 5D). In contrast, DES PND9 epithelial cells had distinct basal region actin staining and there was still not a clear apical actomyosin network as in controls. Despite the differences in apical actin, tight junction formation marked by ZO-1 staining was similar in both treatment groups (Figure 5E). The persistent alterations in both the apical and basal actomyosin networks indicate that alterations in the expression of ECM genes and their cellular receptors in response to DES exposure alters epithelial apical-basal polarity.

Formation of uterine glands occurs postnatally, beginning as invaginations of luminal epithelium into the mesenchyme (15). This type of morphogenetic process involves constriction of apical actomyosin networks to result in epithelial bending, often accompanied by basal actomyosin network relaxation and a partial EMT (13, 42-45). Indeed, mesenchymal markers *Vim* and *Hoxa10* were expressed in control epithelial cells (Figure 5F), albeit at lower levels than in mesenchymal cells, consistent with the possibility that control epithelium was in a partial mesenchymal state. In the epithelium, both of these genes were reduced in DES compared to controls with the greatest reduction observed for *Hoxa10*. Instead, DES epithelium had upregulated markers of terminal epithelial differentiation such as *Sprr2f* and *Alpl* (Figure 5G), consistent with previous observations (9). To gain a better understanding of the potential EMT state of the epithelial cells, we generated a heat map of the relative fold change of well-established epithelial and mesenchymal markers (46). Both control and DES mesenchymal cells had relatively high expression of mesenchymal markers and low expression of epithelial markers while DES epithelial cells had the opposite (Figure 5H). However, control epithelial cells had a mixed picture, with low expression of several epithelial markers and higher expression of some mesenchymal markers. In fact, the control epithelial cells clustered more closely with the mesenchymal cells than with the DES epithelial cells, indicating that these cells were indeed in a partial EMT state.

Three transcription factor families can control entry into EMT: Snail, Twist, and ZEB (47). Of these, we found that only the *Zeb* genes were significantly expressed in PND5 uterine cells (Datasets S5, S6). *Zeb1* and *Zeb2* were downregulated in DES-exposed epithelium, though still highly expressed in mesenchyme (Figure 5I). These findings suggest that DES-induced repression of *Zeb1* and *Zeb2* expression are responsible for the loss of the normal partial EMT that exists in unexposed epithelial cells.

### DES exposure leads to increased chromatin accessibility

To determine the mechanisms underlying the DES effects, we used our single nucleus multiomics data to link chromatin accessibility to RNA expression data from each cell. Using the annotation based on RNA expression, we subset epithelial and mesenchymal cells and examined their chromatin accessibility. Standard dimensional reduction and clustering was based on ATAC data from both epithelial cells and mesenchymal cells, with cells from control and DES-exposed mice integrated together. In epithelium, DES cells remained in an intermediate space between two populations of control cells in the resulting UMAP (Figure 6A-B). In these cells, there were 5,515 peaks gained with DES exposure, and 653 lost (Dataset S12). In mesenchyme, the UMAP of cells based on ATAC data did not separate by exposure (Figure 6C-D). Only 606 peaks were gained with DES exposure, and 303 lost (Dataset S13).

**Figure 6.**
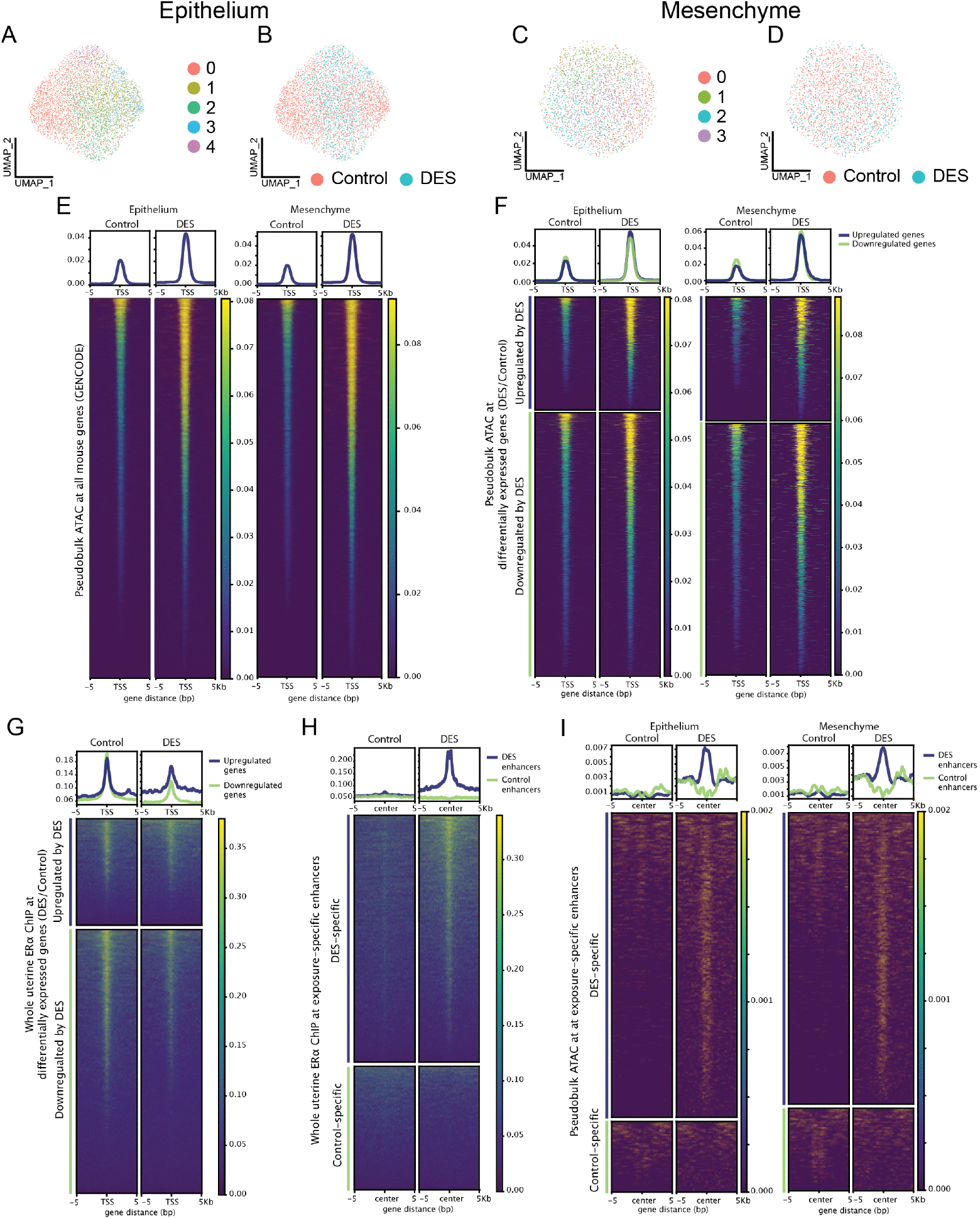
DES promotes chromatin accessibility and alters ERα binding, especially at differential enhancers. A) UMAP of epithelial cells from control or DES-exposed uteri with clustering based on snATACseq data. B) Epithelial ATAC UMAP labeled by exposure. C) UMAP of mesenchymal cells from control or DES-exposed uteri with clustering based on snATACseq data. D) Mesenchymal ATAC UMAP labeled by exposure. E) Metaplots and heatmaps of pseudobulk, normalized ATAC signal at TSS of all GENCODE genes from control or DES-exposed epithelium and mesenchyme. F) Metaplots and heatmaps of ATAC signal at TSS of differentially expressed genes from epithelium and mesenchyme. G) Metaplots and heatmaps of whole uterine ERα ChIP reads at TSS of differentially expressed genes from epithelium and mesenchyme. H) Metaplots and heatmaps of whole uterine ERα ChIP reads at control- or DES-specific enhancers identified in (23). I) Metaplots and heatmaps of pseudobulk ATAC signal from control or DES-exposed epithelium and mesenchyme at control- or DES-specific enhancers (23).

To assess the specificity of changes in accessibility in the two cell types with DES exposure, a pseudobulk, normalized ATAC read density was calculated at the transcription start site (TSS) of all GENCODE genes. DES exposure resulted in a global increase in accessibility in both epithelium and mesenchyme (Figure 6E). We also examined accessibility around the TSS of the subset of differentially expressed genes comparing control and DES-exposed cells (Tables S5-S6). Here as well, there was a global increase in accessibility in DES compared to control cells regardless of the direction of gene expression change (Figure 6F). Because many genes are downregulated in response to DES, this finding highlights the importance of corepressors or other inhibitory elements in regulating gene expression following DES exposure.

DES effects are mediated by ERα, and therefore we wanted to know where ERα binds chromatin in uterine cells at PND5. To do so, we performed ChIPseq on whole PND5 uteri from DES-exposed or unexposed mice. Although this method cannot differentiate ERα binding sites in distinct cell types, whole uteri provide sufficient material for the method, and the results identify ERα binding sites that are used in at least some uterine cells at PND5 rather than using an indirect technique such as a motif analysis. DES exposure resulted in 4,438 unique ERα peaks (of 5,037 total; Dataset S14), compared to only 1,285 peaks unique to controls (of 1,884 total; Dataset S15). Considering only the TSS of genes differentially expressed in DES vs control epithelial and mesenchymal cells (from the multiomics data), there was more enrichment of ERα binding in controls, particularly at genes downregulated by DES (Figure 6G). Genes upregulated by DES also had somewhat reduced ERα binding, suggesting that the promoter regions were not driving gene expression. We next examined ERα binding at the enhancers that have differential H3K27ac marks following DES exposure at PND5 (23). At DES-specific enhancers, there was dramatically increased ERα binding in DES-exposed uteri (Figure 6H), consistent with previous observations that enhancer activation mediates DES effects on the PND5 uterus (23). In contrast, neither treatment group had ERα binding at control-specific enhancers (Figure 6H). To determine how cell type-specific changes in accessibility correlated with the differential enhancers, pseudobulk ATAC read density was mapped at these sites. Accessible chromatin at DES-specific enhancers increased with DES, in both epithelium and mesenchyme, with DES-specific sites more accessible than control-specific sites (Figure 6I). All together, these data demonstrated increased chromatin accessibility co-occurring with ERα binding at DES-specific enhancer sites, but not promoter sites, in DES-exposed cells.

### DES-induced changes in chromatin accessibility at differentially expressed genes correspond to changes in ERα binding sites and chromatin looping

To understand how DES changes the overall chromatin landscape to alter gene expression, we examined specific genes that had differential expression and differential accessibility within the same cell. This process enabled analysis of the correlation (referred to as linkage) between the presence or absence of nearby ATAC peaks and the expression of any gene. Additionally, to assess changes in chromatin architecture, we performed HiC-seq on whole uteri from DES-exposed or unexposed mice at PND5. Although this method was performed using whole tissue, we reasoned that if changes in chromatin looping occurred, these changes were likely to be present in the cell types that were undergoing gene expression changes, whether or not they were also present in cell types that did not have gene expression changes. Of 17,586 total loops found in DES-exposed uteri, and 15,220 in control uteri, 402 loops were gained and 267 lost with DES exposure (Dataset S16). Peaks from bulk ChIP-seq of ERα binding sites and loops from HiC sequencing were overlaid with the aggregate single cell ATAC data and gene-specific linkages to create a picture of how changes in RNA expression could be tied to accessibility, chromatin looping, and ERα binding. Examples follow that illustrate where different combinations of chromatin interactions and ERα binding sites were present at the same locations as cell type-specific changes in ATAC peak strength and RNA expression.

An example of changes in epithelium-specific gene expression linked to changes in chromatin accessibility, ERα binding sites and chromatin looping was provided by branched chain amino acid transaminase 1 (*Bcat1*), an estrogen-regulated gene highly upregulated only in epithelial cells (Figure 7A). The *Bcat1* locus had extensive linkages across a ∼400 kb region that included 5 additional genes, the closest one of which, lymphoid-restricted membrane protein (*Lrmp*), was also upregulated in DES epithelium. Almost all these linkages were positively associated with gene expression and were located at numerous sites of ERα binding and open chromatin largely unique to the DES sample. The single negative linkage was between the *Bcat1* promoter region and a site that had increased ATAC signal in control epithelial cells, suggesting it was a repressor site. There was a very interesting change in chromatin looping between the *Bcat1* promoter and a region ∼80 kb downstream. In controls, the 5′ end of this promoter region loop was connected to a site adjacent to *Lmntd1* that contained an ERα binding site, and the 3′ end of this promoter region loop was looped to a site within the *Casc1* gene that did not have an ERα binding site. In DES cells, the *Bcat1* promoter region loop was switched, with the 5′ end connected to *Casc1* and the 3′ end connected to the site adjacent to *Lmntd1*. At the same time, the chromatin looping site at the *Casc1* locus became extended toward an ERα binding site, and ATAC signal in DES epithelial cells increased in this extended region. The extended region remained looped to the *Bcat1* promoter, likely bringing an active enhancer region to upregulate gene expression. The large, DES-induced epithelium-specific changes in gene expression and chromatin accessibility correlated directly with altered chromatin loops and ERα binding sites at the relevant gene loci and surrounding area.

**Figure 7.**
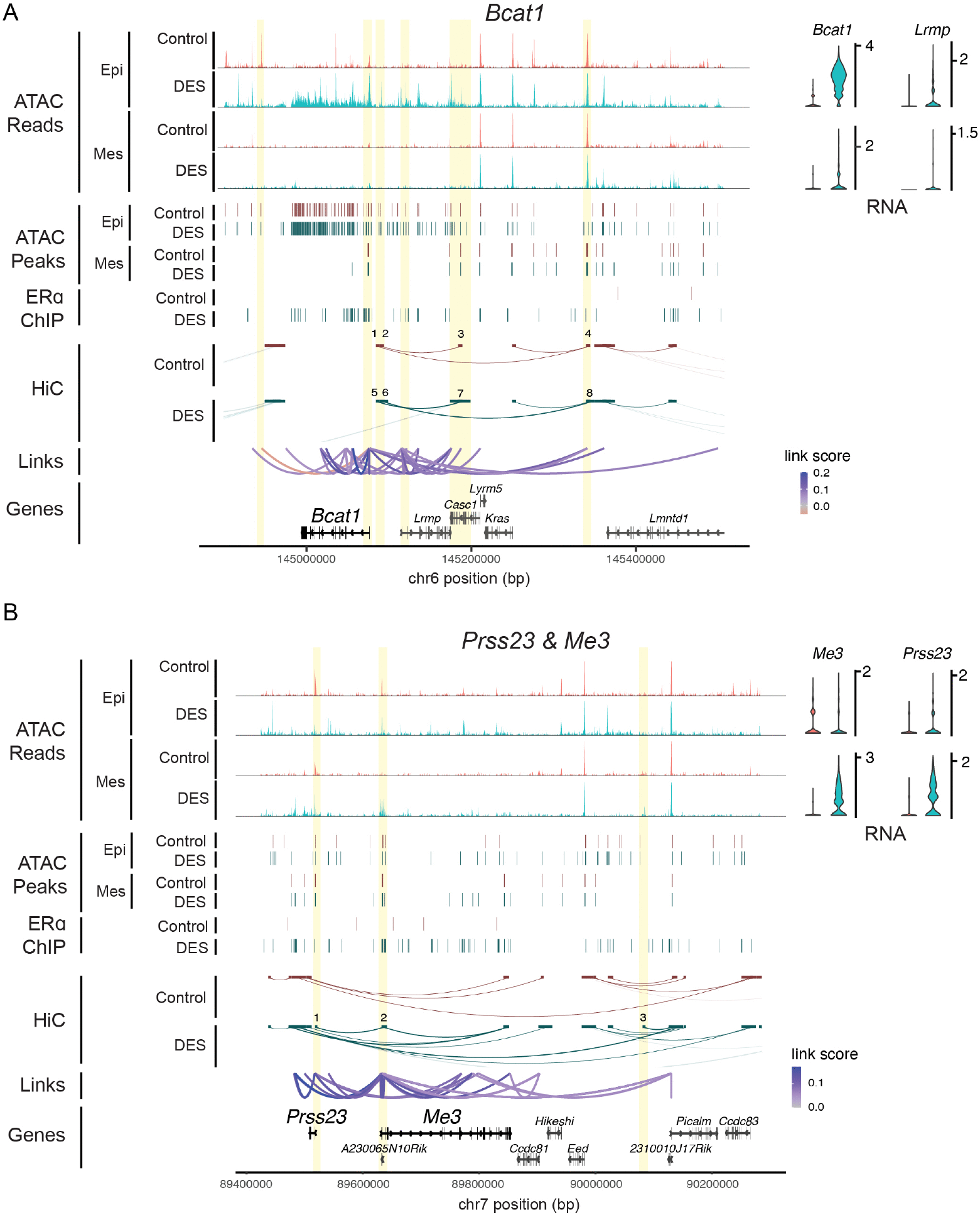
Alterations in chromatin accessibility, looping, and ERα binding co-occur at common sites in the DES-exposed uterine epithelium. A-B) Gene tracks at area surrounding A) *Bcat1* and *Lrmp* or B) *Prss23* and *Me3*, including normalized psuedobulk ATAC signal and peaks in control or DES-exposed uterine epithelial or mesenchymal cells, ERα ChIP peaks from control or DES-exposed whole uterus, HiCseq loops from control or DES-exposed whole uterus, and linkage of RNA expression to open chromatin based on cell-specific ATAC signal. Yellow bars denote areas of interest mentioned in main text. A) 1, control loop at 5’ of promoter region of *Bcat1*; 2, control loop at 3’ of promoter region of *Bcat1*; 3, control loop within *Casc1* with no ERα binding site; 4, control loop near *Lmntd1* with DES-specific ERα binding site; 5, DES loop at 5’ of promoter region of *Bcat1*; 6, DES loop at 3’ of promoter region of *Bcat1*; 7, expanded DES loop within *Casc1* with no ERα binding site; 8, DES loop near *Lmntd1* with DES-specific ERα binding site. B) 1, DES-specific loop at *Prss23* promoter; 2, DES-specific loop at *Me3* promoter; 3, DES-specific loop upstream of *Picalm* with ERα binding site.

Mesenchyme had fewer differentially expressed genes near differentially accessible sites. One area of interest was the chromosome 7 locus that contains serine protease 23 (*Prss23*) and mitochondrial NADP(+)-dependent malic enzyme (*Me3*), which had extensive linkages positively correlated with upregulation by DES in the mesenchyme. At the *Me3* promoter, there was a DES-specific increase in chromatin accessibility in mesenchyme that co-occurred with a DES-specific chromatin loop and ERα binding sites (Figure 7B). This loop was linked to a second DES-specific chromatin loop at the *Prss23* promoter that had an ERα binding site and accessible chromatin, though the accessible chromatin was also present in epithelial cells. A third DES-specific chromatin loop was upstream of the nearby *Picalm* gene at a location with an ERα binding site and minimally increased ATAC signal in DES mesenchyme. This loop was connected directly to the *Picalm* promoter and indirectly to both the *Prss23* and *Me3* promoters. For these genes, expression seemed to be tied to new chromatin looping sites driven by DES-induced ERα binding rather than large changes in chromatin accessibility at the loop sites.

Together, these data support the concept that alterations in cell type-specific gene expression are induced by the combined impacts of changes in chromatin accessibility, chromatin looping, ERα binding, and the association of coactivators and corepressors. The specific mechanisms utilized differ across distinct regions of the genome, but all depend on the presence of ERα in the relevant cell type.

## Discussion

Here we found that developmental exposure to DES promotes premature expression of ERα in uterine epithelium, reduces uterine Wnt signaling, and eliminates the *Lgr5*+ epithelial stem cell population. Intercellular communication is dramatically reorganized, and the developing uterine epithelium loses its normal partial EMT character. Epithelial cell apical-basal polarity changes that under normal conditions likely support glandular development are also disrupted. Mechanistically, DES exposure increases chromatin accessibility and causes cell type-specific changes in gene expression, most prominently in epithelial cells. At least some of the critical gene expression changes can be explained by cell type-specific changes in chromatin accessibility and chromatin looping at enhancer regions in concert with alterations in ERα binding. These findings suggest that ERα-mediated reprogramming of cell differentiation trajectories underlies the long-term phenotypic effects of developmental exposure to estrogenic endocrine disrupting chemicals (Figure 8).

**Figure 8.**
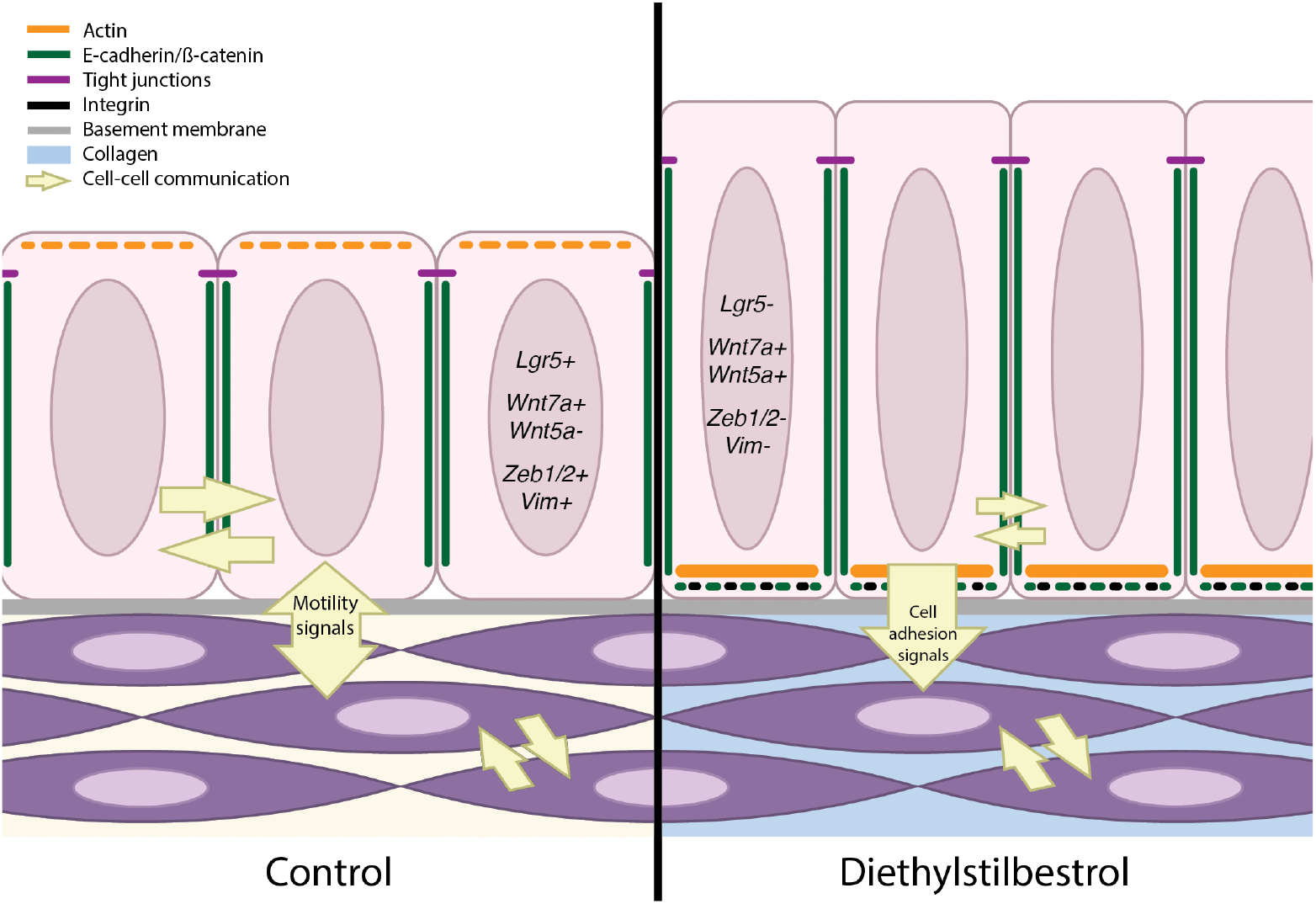
Neonatal DES exposure results in changes to the epithelial stem cell population, cell-cell communication, Wnt signaling, apical-basal polarity, and EMT status. Model of control and DES-exposed uterine epithelium and mesenchyme. Genes in nucleus of epithelial cells reflect key expression differences in response to DES exposure. Yellow arrows indicate direction, amount, and nature of cell-cell signaling. Orange lines within epithelial cells reflect change in actin localization from apical to basal. Green lines represent similar expression of E-cadherin & β-catenin laterally, with aberrant expression punctate in the basal region of the DES-exposed epithelium. Black dots reflect increased integrin expression in the basal region of the DES-exposed epithelium. The change from yellow to blue space between mesenchyme cells reflects the increased collagen expression in DES-exposed cells.

Wnt/β-catenin signaling is a critical mediator of female reproductive tract development based on data from multiple animals including humans (48-51). Estrogens can both activate and inhibit Wnt signaling depending on the cell and tissue context. For example, ERα and β-catenin physically interact to promote transcription of target genes in colon and breast cancer cells, and estradiol promotes translocation of β-catenin to the nucleus in uterine stromal cells (52, 53). Estrogen can also act through a non-genomic mechanism to promote expression of Wnt ligands and frizzled receptors important for uterine epithelial growth (54). However, in the developing uterus it is clear based on the results presented here that the normal developmental balance in Wnt signaling is shifted to full repression by DES exposure. This finding is similar to that observed in a mouse hepatocarcinoma model in which mice treated with an ERα agonist have suppressed Wnt/β-catenin signaling, lower tumor burden and extended survival (55). What causes these differences in the downstream consequences of estrogenic chemical action on Wnt signaling is unknown, but the overall chromatin state and relative expression of coactivators/corepressors in the developing uterus and cancer cells may be key factors.

In addition to its important roles in development, LGR5 is a marker of adult epithelial stem cells in numerous tissues and in cancer stem cells (56). The endometrial epithelium is no exception, as LGR5 is required for both in vivo Wnt signaling-dependent endometrial gland development and ex vivo endometrial epithelial cell self-renewal in the context of organoids (16). Here we extended the previous observation that estradiol treatment inhibits *Lgr5* gene expression in adult uterine tissue (17) to show that DES similarly inhibits *Lgr5* expression in the context of the neonatal, differentiating uterus. This finding likely explains the reduction in uterine gland development observed in adult mice exposed neonatally to either DES or the phytoestrogen genistein (10, 31). However, the presence of some glands in exposed uteri suggests that *Lgr5* expression must resume sufficiently to support their development, even if this does not occur by PND9 (Figure 2F). Alternatively, an additional bipotential epithelial stem cell population could support the limited glandular development (57). Because LGR5 is suppressed by estrogenic chemicals in both developmental and adult settings, long term negative impacts of estrogenic endocrine disruption on stem cells and adult tissue functions could be explained by this mechanism. Furthermore, it raises the possibility that targeting LGR5 using hormonal manipulations could be a useful adjunct to other LGR5-targeting cancer therapies (56).

A partial EMT state is characteristic of epithelia in multiple organs during prenatal organogenesis and of neoplastic cells undergoing invasion or metastasis (58, 59). We uncovered a similar profile of partial EMT in control developing uterine epithelium, suggesting a need for uterine epithelial cells to take on mesenchymal traits at this developmental stage in preparation for endometrial gland budding and extension into the stroma. During morphogenesis of mammary gland branching ducts, the leading epithelial cells undergo partial EMT so that they lose stable epithelial phenotypes and can invade the surrounding stroma (60). This is a tightly regulated process, influenced by Wnt signaling, growth factor signaling, and classical EMT transcription factors such as Snail, Twist, and Zeb family proteins. It is accompanied by expression of matrix metalloproteases that break down ECM material to facilitate epithelial migration into the stroma (61). Here we show that similar mechanisms are likely operating in normal developing endometrium, where the epithelium expresses *Zeb1* and the epithelium and stroma express several matrix metalloproteases. DES exposure pushes the normal partial EMT to a full epithelial state with stable adhesion to the basement membrane supported by a basal actin network, combined with reduced MMP expression and elevated stromal collagens. The combined impact of these changes likely underlies the observed impairment in uterine adenogenesis.

During prenatal female reproductive tract development from the Müllerian duct, cell-cell-communication is vital to control the developmental plasticity of the tissue (62). This remains true during the mouse neonatal period when mesenchymal signals control epithelial differentiation patterns (25). The main ways this communication is regulated are through Wnt signaling and Hox transcription factor-mediated gene expression (62). As reproductive maturity is reached, mesenchymal retinoic acid signaling becomes important for preventing endogenous estrogen-induced transdifferentiation of uterine epithelial cells into stratified squamous epithelium with basal cell differentiation (35). This mechanism, which is ERα-dependent, suggests a need for ongoing mesenchymal-epithelial communication to maintain proper epithelial identity.Restructuring of cell-cell communication following neonatal DES exposure (Figure 3B), compounded by dramatic changes in mesenchyme ECM composition (Figure 3D, 4C), clearly impairs maintenance of the normal epithelial phenotype and likely explains the basal cell metaplasia and squamous metaplasia observed in adult uterine epithelium of mice exposed neonatally to DES or genistein (12).

An important question regarding this work is why the developmental timing is so important in the disruptive effects of neonatal DES exposure. We propose that the critical factor is that DES (or other estrogenic chemicals) induces the expression of ERα in epithelial cells when it is not normally present at sufficient levels to be functional (26), resulting in a loss of the partial EMT characteristics that are required for adenogenesis. The change in ERα functional status causes the epithelial cells to be susceptible to ERα-mediated differentiation signals that are activated in the presence of continued DES exposure over several days, at time points in which the protective effects of retinoic acid signaling are not yet active. These ERα-mediated signals promote global changes in chromatin accessibility and three-dimensional chromatin structure. Furthermore, the DES exposure leads to ECM deposition responses in the mesenchyme that impair cell-cell communication – likely also preventing quick reversal of the abnormal epithelial differentiation phenotypes when DES is discontinued. Later in development, processes such as retinoic acid signaling limit epithelial remodeling by either endogenous or exogenous estrogenic chemicals, protecting the uterine epithelium from long term phenotypic consequences.

During human reproductive tract development, there is no *Esr1* expression in Mullerian duct derivatives until >10 post-conceptional weeks, and likely no presence of functional ERα signaling in the epithelium until sometime after 17 post-conceptional weeks (48). However, in human endometrial organoids derived from 12 and 17 post-conceptional week tissues, estrogenic endocrine disrupting compounds induce *ESR1* and *PGR* expression (48). If our proposal that the timing of ERα expression in the developing female reproductive tract is key for understanding the window of susceptibility to estrogenic endocrine disruptors, these findings suggest that the human female fetus is susceptible at least well into the second trimester. The well documented impact of DES exposure during fetal development on adult female reproductive tract differentiation and function validates this concept (2). Given that uterine adenogenesis is ongoing from the second trimester through puberty (63), this susceptibility may persist. Indeed, neonatal exposure to soy based infant formula, a plant source of estrogenic chemical exposure, increases the incidence and/or severity of common adult human reproductive tract pathophysiological conditions, including uterine fibroids, heavy menstrual bleeding, and endometriosis (64-67). Such pathologies could have their beginnings in the impact of environmental exposures on cell differentiation trajectories during development.

## Materials and Methods

Animal procedures were approved by the National Institute of Environmental Health Sciences (NIEHS) animal care and use committee under protocol 2007-038. Animal care, treatments and tissue collection were performed as described in (11). Tissues were fixed as described in (68). Single-nucleus isolation and sequencing protocol was adapted from (11). ChIP and HiC protocols adapted from (20). Full details regarding analyses and other protocols can be found in Supplemental Methods, with key reagents and software described in Key Reagent table. Code used for analysis is found at https://github.com/DrBrainbridge/PND5-snMultiomics.

## Abbreviations

ATAC: assay for transposase-accessible chromatin
ChIPseq: chromatin immunoprecipitation sequencing
DES: diethylstilbestrol
ECM: extracellular matrix
EMT: epithelial-mesenchymal transition
ERα: estrogen receptor alpha
HiC: chromatin conformation capture
PGR: progesterone receptor
PND: postnatal day
scRNAseq: single cell RNA sequencing
snATACseq: single nucleus ATAC sequencing
snRNAseq: single nucleus RNA sequencing
TSS: transcription start site
ub: ubiquitination
UMAP: uniform manifold approximation and projection

## Acknowledgments

This research was supported by the Intramural Research Program of the National Institutes of Health (NIH), National Institute of Environmental Health Sciences, 1ZIAES102405 (CJW). The contributions of the NIH author(s) are considered Works of the United States Government. The findings and conclusions presented in this paper are those of the author(s) and do not necessarily reflect the views of the NIH or the U.S. Department of Health and Human Services.

