## Supplemental Info for "Neonatal diethylstilbestrol exposure disrupts uterine epithelial apical-basal polarity and partial EMT state"

### Supporting Information Text

#### Data Sharing
Data will be made accessible through NLM BioProject.

#### Supplemental Methods

#### Animal care, DES exposure, and tissue collection
Timed pregnant CD-1 mice were obtained from the in-house breeding colony at NIEHS, housed in a 12 h light:dark cycle, fed NIH-31 diet and water ad libitum. Following delivery, pups were randomly standardized to 10 female pups per dam. Pups were subcutaneously injected with DES (1mg/kg/day; Sigma-Aldrich) or corn oil (vehicle control) for the first five days after birth. At PND5, mice were sacrificed, 4 h after injection. Uteri for single nucleus multiomic sequencing were collected, pooled, fixed in Lomant's reagent (Dithiobis(succinimidyl propionate), DSP; ThermoFisher) and frozen to -70°C in a Mr. Frosty Freezing container (ThermoFisher), following the FixNCut protocol (1). Uteri collected for ChIPseq or HiCseq were frozen immediately after collection on dry ice. Uteri collected for RNAscope, immunohistochemistry, or histology were fixed in 10% neutral buffered formalin.

#### Single nucleus multiomic sequencing
The nucleus extraction protocol was adapted from a previously published protocol for single-cell RNA-sequencing in DES-exposed adult mouse uterus (2). Fixed, frozen uterine tissue was crushed in a dry ice-cooled metal mortar and pestle, and 40-60 mg (uteri of approximately 10 mice per pooled sample) were used for nuclear isolation. Tissue homogenization was performed in Homogenization buffer (composition below) including NP40, digitonin, AlbuMAX bovine serum albumin (BSA), and Protector RNase inhibitor. The suspension was then incubated in OMNI Buffer RSB (composition below) with RNAse inhibitor on ice for 3 min. Nuclei were released by fragmentation in small and large cell dounce homogenizers. Nuclei were filtered through 100 µm and 30 µm CellTrics filters, centrifuged at 4000 rpm for 10 min at 4°C, then resuspended in 10X Nuclei buffer at a concentration of $1x10^6$ cells/mL for library preparation.

<u>OMNI Buffer RSB</u>

| Reagent | Final Concentration |
|---|---|
| Tris-HCl (pH = 7.4) | 10 mM |
| NaCl | 10 mM |
| $MgCl_2$ | 3 mM |
| Tween-20 | 0.1% |
| | Store at 4°C |

<u>Homogenization buffer</u>

| Reagent | Final Concentration |
|---|---|
| NP40 | 0.1% |
| Digitonin | 0.01% |
| BSA | 0.9% BSA |
| RNase inhibitor | 1 U/uL |
| OMNI RSB | Bring to desired volume |

The 10X Genomics Chromium Next GEM Single Cell Multiome ATAC + Gene Expression Library Preparation Kit was used to generate a single nucleus barcoded library of mRNA and transposed DNA from individual nuclei. Single nuclei were counted on a Luna-Fx7 cell counter with AO/PI dyes and examined for quality on a fluorescent microscope. About 16000 nuclei were loaded onto 10X Chip J followed by generating GEMs (Gel Bead-In Emulsions) on a 10X Chromium Controller. In each droplet of GEMs, mRNA is reverse transcribed and open chromatin is transposed with the same cell barcode. The reverse transcribed cDNA and transposed genomic DNA are amplified and separated to construct RNA-seq and ATAC-seq libraries following the instructions. The RNA-seq libraries are sequenced with parameters of 28 nt for read 1 and 90 nt for read 2. The ATAC-seq libraries are sequenced with parameters of 50 nt for read 1 and 49 nt for read 2. Sequencing was done on an Illumina NOVAseq 6000 with sequencing depth of $1x10^8$ read pairs/nucleus for the RNA-seq libraries and $1x10^8$ read pairs/nucleus for the ATAC-seq libraries.

**Single nucleus multiomic analysis**
Most analysis was performed with the Seurat (3) and Signac (4) packages. We performed normalization with SCTransform (5) and in some cases, integrated based on this transformation. Then, we performed dimensional reduction with principal component analysis (PCA) and uniform manifold approximation and projection (UMAP), followed by clustering based on shared nearest neighbor graph. Significantly differential peaks were defined with adjusted p-value < 0.05, average $log_2$ fold change > |1.5|. RNA expression heatmaps were generated using Seurat or pheatmap (6). Ring plots were generated using scplotter (7). To assess cell-cell communication, LIANA was used (8). Pseudobulk ATAC signal tracks were generated after subsetting the CellRanger snATAC BAM file based on the CB (cell barcodes) tag. Heatmaps were generated using DeepTools GenerateHeatmap (9).

**ERα ChIPseq**
Uteri collected and fresh flash frozen from PND5 mice (140 mg of tissue per pooled sample group) were sent to Active Motif for ERα ChIP-seq. ChIP seq reads (75 nt, single-end) were generated by Illumina sequencing (NextSeq 500). Raw single-end ChIP-seq reads (75 bp) were filtered to retain reads with an average Phred quality score >20 and aligned to the mouse reference genome (mm10) using Bowtie v1.1.2 with unique mapping and allowing up to two mismatches per read (-m 1 -v 2) (10). PCR duplicates with identical alignment coordinates were removed using the MarkDuplicates function from Picard Tools. The retained reads were extended to 300 bp for downstream analyses. To normalize sequencing depth across ChIP-seq datasets, reads were downsampled to the same number of uniquely mapped reads per sample. A single pooled input sample for diestrus and estrus was generated to verify antibody specificity but was not used for normalization. For visualization, bedGraph files were converted to bigWig format using bedGraphToBigWig (11). The read coverage was displayed as custom tracks on the IGV Genome Browser, then manually overlayed to match signac coverage plots in Adobe Illustrator.

Peak calling was performed using MACS2 with an adjusted p-value cutoff of <0.0001 (12). Differential regions between a pair of ChIPseq peaks were identified using MEDIPS (Model-based Exploration of DiPloid Sequencing data) with a window size of 250 bp (13). Differential analysis was performed using the medips.meth function with false discovery rate (FDR) adjustment (p.adj = "fdr") and edgeR as the statistical method (diff.method = "edgeR"). Differential regions were defined as genomic intervals exhibiting at least a two-fold difference in mapped read counts and an adjusted p-value ≤ 0.01.

**HiCseq**
Uterine tissues from PND5 control (16 uteri; 54 mg) and DES (5 uteri; 51 mg) were pooled and pulverized on dry ice. Instructions from the Arima Hi-C kit were followed. Sequencing was performed on an Illumina NOVAseq 6000. Raw paired-end Hi-C reads (151 bp) were aligned to the mouse reference genome (mm10) and processed using HiCUP v0.7.1 (14) for read truncation, filtering, and PCR duplicate removal. Only uniquely mapped di-tags that passed quality control and had an intra-chromosomal interaction distance greater than 10 kb were retained for downstream analyses. Chromatin loops were identified using the hiccups algorithm implemented in Juicer v1.8.9 with default parameters (15). The classification of common and differential loops was based on a loop's presence following DES exposure and in controls.

**Immunohistochemistry**

| Target | Staining technique | Epitope retrieval | Non-specific | Primary antibody | Secondary antibody |
|---|---|---|---|---|---|

| | | | site block | | |
|---|---|---|---|---|---|
| Laminin | Avidin-biotin-peroxidase | Proteinase K 5 min at room temperature | 10% normal donkey serum | Rabbit monoclonal laminin antibody (1.0 mg/ml) at a 1:7500 dilution or concentration matched normal rabbit IgG for 1 hour at RT | Biotinylated donkey anti-rabbit IgG at a dilution of 1:500 for 30 min at RT |
| E-cadherin | Avidin-biotin-peroxidase | Heat induced in the Decloaker® pressure chamber for 15 min at 110°C in citrate buffer (pH 6.0) | 10% normal horse serum | Mouse monoclonal E-cadherin antibody (0.25 mg/ml) or a concentration matched normal mouse IgG2a at a 1:1000 dilution for 1 hour at RT | Biotinylated horse anti-mouse IgG at a dilution of 1:1000 |
| B-catenin | HRP-polymer | Heat induced in the Decloaker® pressure chamber for 15 min at 110°C in citrate buffer (pH 6.0) | NA | β-catenin antibody at 1:2500 dilution for 30 min at RT | Rabbit on Rodent HRP Polymer for 15 min at room temperature |
| ZO-1 | Avidin-biotin-peroxidase | Heat induced in the Decloaker® pressure chamber for 15 min at 110°C in in EDTA buffer solution (pH 8.5) | 10% normal horse serum | Mouse monoclonal ZO-1 antibody (1.0 mg/ml) at a 1:200 dilution or concentration matched normal mouse IgG1 for 1 hour at RT | Biotinylated horse anti-mouse IgG at a dilution of 1:1000 |

Immunohistochemical staining was performed using the techniques specified in the table above. Formalin-fixed, paraffin-embedded whole uterus tissue sections were deparaffinized in xylene and rehydrated through graded ethanol. Enzyme-induced epitope retrieval was performed. Endogenous peroxidase blocking was done by immersing the sections in 3% $H_2O_2$ for 15 min. Non-specific sites were blocked by incubating slides for 20 min in serum. Endogenous biotin and avidin binding sites were blocked using an avidin-biotin blocking kit. Slides were incubated with primary antibody, then secondary antibody. Labeling incubation was done with the Vectastain RTU Kit Label for 30 min at RT. The antigen-antibody complex was visualized using 3-diaminobenzidine (DAB) chromogen for 6 min. Slides were counterstained with modified Harris hematoxylin, dehydrated through graded ethanol and coverslipped. Images were scanned using a Hamamatsu NanoZoomer S360 Digital Slide Scanner. Gamma correction was performed in NDP.view 2 (Hamamatsu) to level 1.8 unless otherwise stated in figure legend.

**RNAscope**
Formalin-fixed, paraffin-embedded whole uterine horns from control and DES-exposed mice were deparaffinized in xylene, rehydrated, blocked, and probed according to the RNAscope Multiplex Fluorescent Reagent Kit v2 instructions (Document number UM 323100, Advanced Cell Diagnostics, Inc). Opal 570 was used with the *Lgr5* probe on channel 1 (Advanced Cell Diagnostics, Inc), and Opal 690 on the *Esr1* probe on channel 2. Tissues were imaged with a Zeiss LSM 980 inverted confocal microscope with AiryScan.

**Histology**

Whole uterine horns from control and DES-exposed mice were fixed in 10% neutral buffered formalin for approximately 48 h, moved to 70% ethanol, processed and embedded in paraffin blocks and sectioned at 5 μm thickness. For actin staining, following deparaffinization in xylene and ethanol, tissues were permeabilized with 0.1% Triton X-100 for 10 min. They were then incubated in 1:1000 phalloidin-rhodamine, washed, and counterstained with DAPI. Finally, tissues were imaged with a Zeiss LSM 980 inverted confocal microscope with AiryScan. Masson's Trichrome staining was performed according to standard procedures (16). Images were scanned using a Hamamatsu NanoZoomer S360 Digital Slide Scanner (Hamamatsu Corporation, Hamamatsu City, Japan).

**Key Reagent Table**

| REAGENT or RESOURCE | SOURCE | IDENTIFIER |
|---|---|---|
| **Antibodies** | | |
| Laminin Polyclonal Antibody | Thermo Fisher Scientific | Cat# PA1-16730, RRID:AB_2133633 |
| E-cadherin Monoclonal Antibody | BD Biosciences | Cat# 610182, RRID:AB_397581 |
| Beta Catenin antibody | Abcam | Cat# ab16051, RRID:AB_443301 |
| ZO-1 Recombinant Mouse Monoclonal Antibody (ZO1-1A12) | Thermo Fisher Scientific | Cat# 740002M, RRID:AB_3074173 |
| ERα antibody | Santa Cruz | Cat# sc-543 |
| Normal rabbit IgG | Abcam | N/A |
| Normal mouse IgG2a | BD Pharmingen | N/A |
| Normal mouse IgG1 | BD Pharmingen | N/A |
| Biotinylated donkey anti-rabbit IgG antibody | Jackson ImmunoResearch | 1:500 dilution |
| Biotinylated horse anti-mouse IgG antibody | Vector Laboratories | 1:1000 dilution |
| Rabbit on Rodent HRP Polymer | Biocare | Cat# RMR622, lot 100923A |
| **Chemicals, peptides, and recombinant proteins** | | |
| RNAscope™ Multiplex Fluorescent Detection Kit v2 | ACD | Cat# 323110 |
| RNAscope pretreatment reagents (Target retrieval, Pretreatment 2) | ACD | Cat# 322000 |
| RNAscope pretreatment reagents (Hydrogen peroxide, Protease Plus, Protease III, and Protease IV) | ACD | Cat# 322381 |
| RNAscope Wash buffer reagents | ACD | Cat# 310091 |
| RNAscope® Probe - Mm-Lgr5 | ACD | Cat# 312171 |
| RNAscope® Probe - Mm-Esr1-C2 | ACD | Cat# 478201-C2 |
| Opal 570 Reagent Pack | Akoya Biosciences | FP1488001KT |
| Opal 690 Reagent Pack | Akoya Biosciences | FP1497001KT |
| Phalloidin-rhodamine | Thermo Fisher Scientific | Cat# R415 |
| Bouin's Solution | Lab Chem Inc. | Cat# LC11790-1 |
| Hematoxylin crystals | Chroma-Gesellschaft | Cat# 5B 535 |
| Ferric Chloride, 29% aqueous | Thermo Fisher Scientific | Cat# 188-100 |
| Hydrochloric acid, concentrated | Thermo Fisher Scientific | Cat# A508-500 |

| | | |
|---|---|---|
| Biebrich scarlet | Rowley Biochemical | CAS: 4196-99-0 |
| Acid fuchsin | Chroma-Gesellschaft | Cat# 1B 525 |
| Phosphomolybdic acid | Thermo Fisher Scientific | Cat# A237-100 |
| Phosphotungstic acid | Thermo Fisher Scientific | Cat# A248-100 |
| Aniline blue | Thermo Fisher Scientific | Cat# A-967 |
| Glacial acetic acid | Thermo Fisher Scientific | Cat# A35-500 |
| Light green, SF yellowish | Rowley Biochemical | CAS: 5141-20-8 |
| Diethylstilbestrol (DES) | Sigma-Aldrich | CAS: 56-53-1 |
| Corn oil (vehicle control) | Spectrum Chemical | CAS: 8001-30-7 |
| Dithiobis(succinimidyl propionate) (DSP), Lomant's reagent | ThermoFisher | Cat# 22586 |
| Protector RNase inhibitor | Sigma-Aldrich | SKU 3335402001 |
| Proteinase K | Leica Biosystems | Cat# RE7160-K |
| 3-diaminobenzidine (DAB) chromogen | Dako | N/A |
| Normal donkey serum | Jackson ImmunoResearch | N/A |
| Normal horse serum | Jackson ImmunoResearch | N/A |
| Citrate buffer (pH 6.0) | Biocare Medical | N/A |
| EDTA buffer solution (pH 8.5) | Biocare Medical | N/A |
| **Kits** | | |
| 10X Genomics Chromium Next GEM Single Cell Multiome ATAC + Gene Expression Library Preparation Kit | 10X Genomics | Cat# 1000285 |
| Vectastain RTU Kit Label | Vector Laboratories | N/A |
| Avidin-biotin blocking kit | Vector Laboratories | N/A |
| Arima Hi-C kit | Arima | N/A |
| **Experimental models: Organisms/strains** | | |
| CD-1 mice (timed pregnant) | NIEHS in-house breeding colony | N/A |
| **Oligonucleotides** | | |
| See RNAscope probes above | | |
| **Sequence-based reagents** | | |
| See RNAscope probes above | | |
| **Software and algorithms** | | |
| R | R Project | v4.3.2; https://www.r-project.org/ |
| Python | Python Software Foundation | v3.9.6 & v3.11.6; https://www.python.org/ |
| numpy | Harris et al (17) | v1.26.4 |
| pandas | The pandas development team (18) | v2.1.1 |
| scipy | Virtanen et al (19) | v1.15.1 |

| | | |
|---|---|---|
| anndata | Virshup et al (20) | v0.10.9 |
| matplotlib | Hunter (21) | v3.9.4 |
| scanpy | Wolf, et al (22) | v1.10.3 |
| seaborn | Waskom (23) | v0.13.2 |
| NDP.view2Plus | Hamamatsu | https://www.hamamatsu.com/ |
| BedGraphToBigWig | Kent et al (11) | v2.10 |
| samtools | Li et al (24) | v1.18 |
| bedtools | Quinlan & Hall (25) | v2.31.1 |
| deepTools | Ramírez et al (9) | v3.5.1 |
| pheatmap | Kolde (6) | v1.0.12 |
| UpSetR | Conway et al (26) | v1.4.0 |
| MEDIPS | Lienhard et al (13) | v1.54.0 |
| Rsamtools | Morgan et al (27) | v2.18.0 |
| scplotter | Wang (7) | v0.1.1 |
| readxl | Wickham & Bryan (28) | v1.4.3 |
| liana | Dimitrov et al (8) | v0.1.13 |
| BSgenome.Mmusculus.UCSC.mm10 | Team TBD (29) | v1.4.3 |
| BSgenome | Pagès (30) | v1.70.2 |
| BiocIO | Morgan et al (31) | v1.12.0 |
| Biostrings | Pagès et al (32) | v2.70.3 |
| XVector | Pagès & Aboyoun (33) | v0.42.0 |
| EnsDb.Mmusculus.v79 | Rainer (34) | v2.99.0 |
| ensembldb | Rainer et al (35) | v2.26.0 |
| AnnotationFilter | Morgan & Rainer (36) | v1.26.0 |
| GenomicFeatures | Lawrence et al (37) | v1.54.4 |
| SparseM | Koenker & Ng (38) | v1.84-2 |
| AnnotationDbi | Pagès et al (39) | v1.64.1 |
| graph | Gentleman et al (40) | v1.80.0 |
| limma | Ritchie et al (41) | v3.58.1 |
| stringr | Wickham (42) | v1.5.1 |
| dplyr | Wickham et al (43) | v1.1.4 |
| readr | Wickham et al (44) | v2.1.5 |
| tibble | Müller & Wickham (45) | v3.2.1 |
| tidyverse | Wickham et al (45) | v2.0.0 |
| scMerge | Lin et al (46) | v1.18.0 |
| scran | Lun et al (47) | v1.30.2 |
| scuttle | McCarthy et al (48) | v1.12.0 |
| MAST | Finak et al (49) | v1.28.0 |
| SingleCellExperiment | Amezquita et al (50) | v1.24.0 |
| SummarizedExperiment | Morgan et al (51) | v1.32.0 |
| Biobase | Huber et al (52) | v2.62.0 |
| GenomicRanges | Lawrence et al (37) | v1.54.1 |
| GenomeInfoDb | Arora et al (53) | v1.38.8 |

| | | |
|---|---|---|
| IRanges | Lawrence et al (37) | v2.36.0 |
| S4Vectors | Pagès et al (54) | v0.40.2 |
| BiocGenerics | Huber et al (52) | v0.48.1 |
| MatrixGenerics | Ahlmann-Eltze et al (55) | v1.14.0 |
| matrixStats | Bengtsson (56) | v1.5.0 |
| tidyr | Wickham et al (57) | v1.3.1 |
| patchwork | Pedersen (58) | v1.2.0 |
| ggrepel | Slowikowski (59) | v0.9.6 |
| ggplot2 | Wickham (60) | v3.5.2 |
| irlba | Baglama & Reichel (61) | v2.3.5.1 |
| Signac | Stuart et al (4) | v1.14.0 |
| SeuratDisk | | v0.0.0.9021 |
| Seurat | Hao et al (3) | v5.3.0 |
| SeuratObject | Satija et al (62) | v5.1.0 |
| sp | Pebesma & Bivand (63)<br>Bivand et al (64) | v2.2-0 |
| Bowtie | Langmead et al (10) | v1.1.2 |
| MACS | Zhang et al (12) | |
| HOMER | Heinz et al (65) | v4.5 |
| IGV genome browser | Robinson et al (66) | v2.18.4 |
| HiCUP | Wingett (14) | v0.7.1 |
| Juicer | Durand et al (67) | v1.8.9 |
| SCTransform | Choudhary & Satija (5) | |
| Claude | Anthropic | https://claude.ai |
| **Other** | | |
| NIH-31 diet | N/A | N/A |
| 100 µm CellTrics filters | Fisher Scientific | N/A |
| 30 µm CellTrics filters | Fisher Scientific | N/A |
| Mr. Frosty Freezing container | ThermoFisher | N/A |
| 10X Chip J | 10X Genomics | N/A |
| Small cell dounce homogenizer | Kontes Glass Co. | N/A |
| Large cell dounce homogenizer | Kontes Glass Co. | N/A |
| 10X Chromium Controller | 10X Genomics | N/A |
| Illumina NOVAseq 6000 | Illumina | N/A |
| Illumina NextSeq 500 | Illumina | N/A |
| Hamamatsu NanoZoomer S360 Digital Slide Scanner | Hamamatsu Corporation | N/A |
| Zeiss LSM 980 inverted confocal microscope with AiryScan | Zeiss | N/A |
| Decloaker® pressure chamber | Biocare Medical | N/A |
| Luna-Fx7 cell counter with AO/PI dyes | Logos Biosystems | N/A |

### Figures

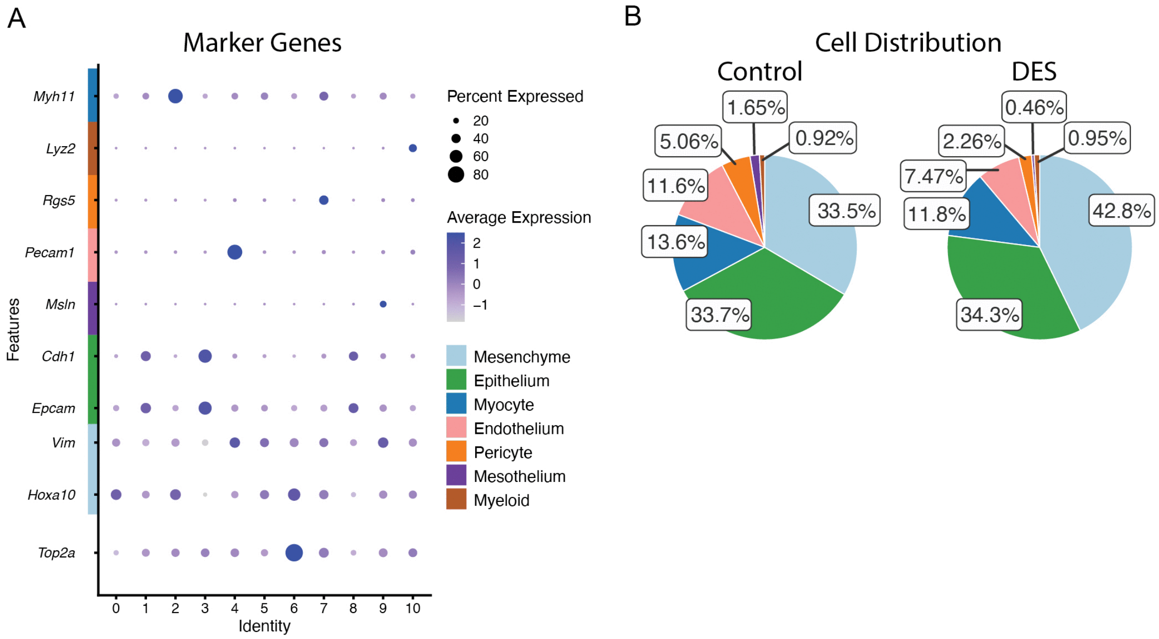


**Fig. S1. Manual annotation of integrated cells from control and DES-exposed uteri.** A) Dot plot of markers for proliferative cells (*Top2a*), mesenchyme (*Vim* & *Hoxa10*), epithelium (*Epcam* & *Cdh1*), mesothelium (*Msln*), endothelium (*Pecam1*), pericyte (*Rgs5*), myeloid (*Lyz2*), and myocyte (*Myh11*). B) Relative composition of control (left) and DES-exposed (right) uterine cells as annotated in Figure 1B.

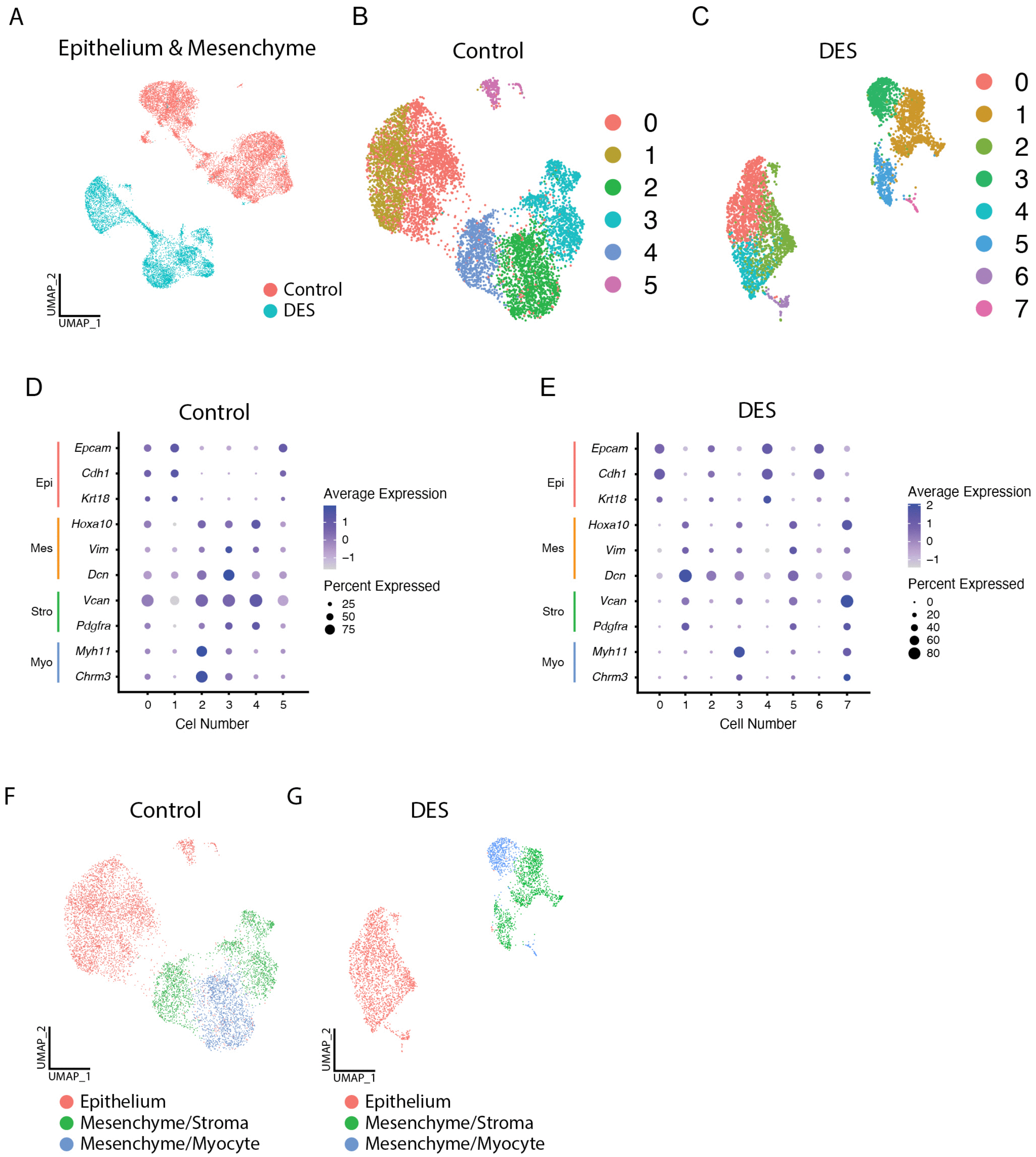


**Fig. S2. Manual annotation of nonintegrated cells from control and DES-exposed uterine epithelium and mesenchyme.** A) UMAP of integrated epithelium, mesenchyme, and myocytes from both control and DES-exposed uterus, labeled by exposure. B) UMAP of integrated control uterine epithelium, mesenchyme, and myocytes labeled by cluster. C) UMAP of integrated DES-exposed uterine epithelium, mesenchyme, and myocytes labeled by cluster. D) Dot plot of expression of markers for myocyte (*Myh11*), stroma (*Dcn*, *Vcan,* & *Pdgfra*), epithelium (*Krt18* & *Epcam*), and mesenchyme (*Vim* & *Hoxa10*) in control uterine epithelium and mesenchyme. E) Dot plot of expression of markers in DES-exposed epithelium and mesenchyme. F-G) Annotated UMAPs of integrated control (F) and DES-exposed (G) uterine epithelium, mesenchyme with additional stromal markers, and mesenchyme with additional myocyte markers.

**Dataset S1 (separate file).** Differentially expressed genes per cluster (whole uterus)

**Dataset S2 (separate file).** Top 25 marker genes per cell type and exposure (whole uterus)

**Dataset S3 (separate file).** Differentially expressed genes per cluster (Control; epithelial and mesenchymal subset)

**Dataset S4 (separate file).** Differentially expressed genes per cluster (DES; epithelial and mesenchymal subset)

**Dataset S5 (separate file).** Epithelial differentially expressed genes (DES/Control)

**Dataset S6 (separate file).** Mesenchymal differentially expressed genes (DES/Control)

**Dataset S7 (separate file).** Liana top 50 ligand-receptor pairs as shown in Figure 2B

**Dataset S8 (separate file).** Liana top 50 ligand-receptor pairs, expression and specificity (Control epithelium & mesenchyme)

**Dataset S9 (separate file).** Liana top 50 ligand-receptor pairs, expression and specificity (DES epithelium & mesenchyme)

**Dataset S10 (separate file).** Liana all ligand-receptor pairs (Control epithelium & mesenchyme)

**Dataset S11 (separate file).** Liana all ligand-receptor pairs (DES epithelium & mesenchyme)

**Dataset S12 (separate file).** Differential ATAC peaks (DES/control epithelium)

**Dataset S13 (separate file).** Differential ATAC peaks (DES/control mesenchyme)

**Dataset S14 (separate file).** ERα ChIPseq peaks (DES whole uterus)

**Dataset S15 (separate file).** ERα ChIPseq peaks (Control whole uterus)

**Dataset S16 (separate file).** Differential HiC (DES/Control whole uterus)
